# Frontal theta phase modulates asymmetric posterior neural mechanisms of spatial attention

**DOI:** 10.64898/2026.03.25.714015

**Authors:** Megan Darrell, Theo Vanneau, Chloe Brittenham, John J. Foxe, Sophie Molholm

**Affiliations:** Medical Scientist Training Program, Albert Einstein College of Medicine, Bronx, New York, USA; Dominic P. Purpura Department of Neuroscience, Albert Einstein College of Medicine, Bronx, New York, USA; The Cognitive Neurophysiology Laboratory, Department of Pediatrics, Albert Einstein College of Medicine, Bronx, New York, USA; The Frederick J. and Marion A. Schindler Cognitive Neurophysiology Laboratory, The Ernest J. Del Monde Institute for Neuroscience, University of Rochester School of Medicine and Dentistry, Rochester, New York, USA

## Abstract

Selective attention enables prioritization of behaviorally relevant information through coordinated control of neural excitability. Although theta-band (3–7 Hz) rhythms are implicated in top-down attentional sampling in non-human primates, how intrinsic theta phase organizes sensory gain and behavior in humans, and whether this control operates symmetrically across hemispheres, remains unknown. We recorded electroencephalography (EEG) and pupillometry in typically developing human participants (n = 21; 14.7 ± 3.8 YO) performing a covert spatial attention task. Behaviorally, participants responded faster during leftward relative to rightward attention. This behavioral asymmetry was paralleled in the neural data: anticipatory modulation of parieto-occipital alpha and beta power emerged selectively during leftward attention, whereas rightward attention did not recruit comparable posterior oscillatory processes.

Mechanistically, ipsilateral fronto-central theta phase emerged as a potential driver of this asymmetry. Intrinsic theta phase predicted trial-by-trial reaction time (RT) in a cue-direction–specific manner. During leftward attention, 3-Hz theta-phase over left fronto-central cortex modulated behavior and was significantly coupled to coordinated posterior alpha-band activity. In contrast, 6–7-Hz theta-phase over right fronto-central cortex modulated behavior during rightward attention but showed no relationship with alpha or beta modulation; instead, it modulated early sensory gain, indexed by P1 amplitude. Consistent with these distinct architectures, RT was jointly predicted by lower pre-stimulus alpha power and higher P1 amplitude over the attended hemisphere during leftward attention, whereas only P1 amplitude predicted performance during rightward attention. Resting-state alpha power did not differ across hemispheres, indicating that these effects were task-evoked rather than baseline spectral differences. Critically, older participants, who demonstrated enhanced behavioral performance, also exhibited a larger hemispheric asymmetry. Together, these findings reveal developmentally emerging, direction-specific neural control dynamics underlying human spatial attention.

**Significance Statement:** Spatial attention is often assumed to rely on symmetric neural mechanisms across left and right space. Using EEG in typically developing children and adolescents, we show that intrinsic theta rhythms organize attention through direction-specific control architectures. Leftward attention engages slower frontal theta (3-Hz) that coordinates posterior alpha and beta activity, consistent with oscillatory sensory gating. Rightward attention instead relies on faster theta (6–7-Hz) that modulates early sensory responses without coordinated alpha dynamics. These asymmetric mechanisms occur despite lack of hemispheric differences in resting alpha activity, indicating that they emerge during active control rather than reflecting baseline biases. These findings reveal that human attentional sampling is rhythmically organized but fundamentally asymmetric across space.

## Introduction

Imagine traveling through a crowded city, amid flashing billboards, traffic noise, and streams of pedestrians. In a world flooded with competing sensory information, we harness selective attention to focus on certain stimuli at the exclusion of others (James 1890), which fundamentally shapes how we navigate the world, learn (Keller, Davidesco et al. 2020), make decisions (Krajbich, Armel et al. 2010, Krajbich and Rangel 2011, Perkovic, Schoemann et al. 2023), and form social connections (Frischen, Bayliss et al. 2007, Nummenmaa and Calder 2009).

Desimone and Duncan (1995) proposed a behavioral and neural framework for the selective attention process grounded in two core principles: (1) the brain has limited capacity to process information, and therefore (2) stimuli must compete for neural representation (Desimone and Duncan 1995). Selective attention thus serves to bias this competition in favor of behaviorally relevant targets, guided by current goals and expectations. Within this competitive architecture, spatial attention emerges as a key mechanism for resolving representational conflict across the visual field. It engages large-scale, hemispheric control networks, including parietal cortex (Petersen, Corbetta et al. 1994, Townsend and Courchesne 1994, Foxe, McCourt et al. 2003, Thakral and Slotnick 2009, Szczepanski, Konen et al. 2010), the superior colliculus ((Hu and Dan 2022); see (Krauzlis, Lovejoy et al. 2013) for review), and thalamic (pulvinar) circuits (Petersen, Robinson et al. 1987, Zhou, Schafer et al. 2016), that bias sensory processing toward relevant locations (Colby 1991). Critically, these dynamics unfold over time and across networks (Miller and Buschman 2013), making electroencephalography (EEG) well-suited to capture the temporal evolution and hemispheric coordination of attentional selection in humans.

A growing body of evidence indicates that these distributed control mechanisms are coordinated through neural oscillations, reflecting synchronized neural firing (Petersen, Robinson et al. 1987, Zhou, Schafer et al. 2016), which regulate when and where sensory information is prioritized or suppressed. Among these neuro-oscillations, theta-band activity (∼3–7 Hz) has emerged as a central mechanism of top-down attentional control. A prominent framework, the rhythmic theory of attention, posits that the phase of theta oscillations determines whether the system occupies (i) externally oriented (“sampling”) or (ii) internally oriented (“gating”) periods (Fiebelkorn and Kastner 2019), thereby directly governing attentional dynamics. This temporal organization produces fluctuations in perceptual sensitivity that dictate when sensory information is preferentially processed versus withheld. Consistent with this account, non-human primate studies show that target detection varies rhythmically at 3-6 Hz, suggesting that spatial attention operates in a rhythmic, rather than continuous, manner (Landau and Fries 2012, Fiebelkorn, Saalmann et al. 2013, Fiebelkorn, Snyder et al. 2013, Dugue, Marque et al. 2015, Fiebelkorn, Pinsk et al. 2018).

Importantly, these theta-phase–dependent fluctuations in behavior appear to be mediated by coordinated interactions with faster frequency bands. Rather than acting in isolation, theta phase is associated with systematic fluctuations in alpha (∼7–13-Hz), beta (∼13–25 Hz) and gamma (>30-Hz) power, yielding alternating states of enhanced sensory sampling (low alpha/beta, high gamma) and sensory suppression (high alpha/beta, low gamma) (Klimesch, Sauseng et al. 2007, Klimesch, Sauseng et al. 2007, Romei, Gross et al. 2010, Hanslmayr, Gross et al. 2011, Wilson and Foxe 2020, Han, Brincat et al. 2026).

Accordingly, “suboptimal” theta phases, which are thought to reflect an internally oriented gating state associated with reduced perceptual sensitivity, are accompanied by elevated alpha and beta power. Alpha-band activity in particular, the dominant intrinsic rhythm of the human brain ((Berger 1929); see (E Basar 1996, Basar, Schurmann et al. 1997) for review), is widely thought to suppress sensory processing by modulating cortical excitability prior to stimulus onset, although the precise mechanisms remain debated (Foxe and Snyder 2011, Bonnefond and Jensen 2025). Traditionally characterized as inhibitory, increased anticipatory alpha power over visual cortical regions has been associated with suppression of unattended spatial locations (Worden, Foxe et al. 2000, Kelly, Gomez-Ramirez et al. 2009), distractor visual stimuli during inter-sensory selective attention tasks (Foxe, Simpson et al. 1998, Fu, Foxe et al. 2001, Gomez-Ramirez, Kelly et al. 2011), and task-irrelevant visual features (e.g., color or motion (Snyder and Foxe 2010)). Critically, alpha-band power indices of cortical suppression of distractors have been related to performance on the primary task (Thut, Nietzel et al. 2006, Kelly, Gomez-Ramirez et al. 2009, Murphy, Foxe et al. 2014). Together, these findings suggest that alpha increases during suboptimal theta phases may mediate the cortical suppression that characterizes this internally oriented state. Within this framework, these alpha-mediated inhibitory processes can be understood as spatially specific mechanisms that are temporally coordinated by the phase of ongoing theta oscillations.

Conversely, “optimal” theta phase corresponds to externally oriented, or “sampling,” periods marked by heightened perceptual sensitivity. During these “sampling” windows, reduced alpha and beta power is thought to release inhibitory gating, increasing cortical excitability. This permissive state is accompanied by transient bursts of high-frequency (>30-Hz) gamma activity (Helfrich, Fiebelkorn et al. 2018), which is widely interpreted as a proxy for cortical excitability and sensory enhancement in humans (Gray, Konig et al. 1989, Gray and Singer 1989, Landau, Schreyer et al. 2015, Rich and Wallis 2017, Watson, Ding et al. 2018). Elevated gamma during these phases has been linked to stronger sensory responses (Gruber, Muller et al. 1999) and faster behavioral performance (Frund, Busch et al. 2007), consistent with an externally oriented sampling state. This interpretation is further supported by extensive evidence that theta–gamma coupling temporally coordinates periods of sampling across a range of cognitive demands (Canolty, Edwards et al. 2006, Tort, Komorowski et al. 2009, Szczepanski, Crone et al. 2014). Similarly, early thalamocortical models propose that tonic firing supports active sensory relay, with gamma oscillations facilitating information routing between cortical regions during attention and active processing (Guillery 2001, Sherman 2001).

Together, this model positions theta as a master control rhythm that periodically modulates alpha/beta-mediated inhibition and gamma-mediated excitation in a regionally specific manner, effectively managing sensory gain to specific spatial locations. By rhythmically alternating between suppression and enhancement, theta orchestrates when the brain samples the environment versus when it withholds processing, enabling efficient, temporally and spatially structured attention.

At the level of sensory cortex, these oscillatory state changes are reflected in early visual-evoked potentials, particularly the P1 component. The P1, a positive deflection over parieto-occipital electrodes approximately 80–130 ms after stimulus onset (Luck, Heinze et al. 1990, Heinze, Mangun et al. 1994, Wijers, Lange et al. 1997, Woldorff, Fox et al. 1997), is widely considered an index of early sensory gain and attentional amplification to prioritized stimuli in extrastriate visual cortex (Awh, Anllo-Vento et al. 2000, Frey, Schmid et al. 2014). Notably, P1 amplitude and latency covary with anticipatory alpha power (Brandt and Jansen 1991, Brandt, Jansen et al. 1991, Gruber, Muller et al. 1999, Fellinger, Klimesch et al. 2011), such that lower alpha states are associated with larger and earlier P1 responses, linking oscillatory modulation of cortical excitability to the strength and timing of early sensory processing. Despite an established relationship between alpha and P1, how early sensory gain mechanisms are temporally organized by slower theta rhythms remains unclear.

Moreover, despite growing behavioral and neural evidence implicating theta-phase as a driver of attentional control through oscillatory sensory-gating mechanisms, the underlying implementation of these dynamics in humans remains poorly understood. Much of the mechanistic evidence to date derives from non-human primate electrophysiology, with comparatively limited direct characterization in humans, and virtually no work examining how these control processes emerge or mature across development. Furthermore, prior studies have largely relied on external perturbations—such as brief flashes of light (Landau and Fries 2012, Landau, Schreyer et al. 2015, Lo, Ke et al. 2025), transcranial magnetic stimulation (Gordon, Belardinelli et al. 2022, Trajkovic, Veniero et al. 2025), or systematic manipulation of cue–target interstimulus intervals (Fiebelkorn, Saalmann et al. 2013, Fiebelkorn, Pinsk et al. 2018, Helfrich, Fiebelkorn et al. 2018, Plochl, Fiebelkorn et al. 2022, Cavanah and Fiebelkorn 2025, Han, Brincat et al. 2026)—to impose or align theta phase. As a result, the role of intrinsic, ongoing theta dynamics, and their trial-by-trial variability, in shaping behavioral performance and neural dynamics, remains largely uncharacterized.

It is also unknown whether these oscillatory control mechanisms operate symmetrically across space. A substantial behavioral literature demonstrates a reliable leftward attentional bias (pseudoneglect) in healthy individuals (Bradshaw, Nathan et al. 1987, McCourt and Jewell 1999, Orr and Nicholls 2005, Charles, Sahraie et al. 2007, Sosa, Teder-Salejarvi et al. 2010), consistent with right-hemisphere dominance for spatial attention (Heilman and Van Den Abell 1980, Foxe, McCourt et al. 2003, Spagna, Kim et al. 2020). Converging clinical evidence further supports this asymmetry, as patients with right-sided parietal lesions experience profound hemi-spatial neglect (Stone, Halligan et al. 1993, Heilman, Valenstein et al. 2000, Bartolomeo and Chokron 2002, Chokron 2003, Corbetta and Shulman 2011). Likewise, functional neuroimaging has revealed that leftward and rightward attention recruit distinct large-scale networks, with leftward attention engaging stronger bilateral fronto-parietal control and rightward attention relying primarily on more limited, contralateral recruitment (Siman-Tov, Mendelsohn et al. 2007). Despite prominent evidence for asymmetric attentional control, most EEG studies of spatial attention collapse across cue direction, implicitly assuming equivalent neural mechanisms for leftward and rightward attention. Consequently, whether theta-mediated sensory gating and early sensory gain control differ as a function of attentional direction has never been directly tested.

Here, we provide the first developmental characterization of how intrinsic frontal theta phase organizes higher frequency neuro-oscillatory activity, early sensory responses, and behavior in humans using EEG during a spatial attention task. Importantly, this investigation is embedded within a multimodal framework that simultaneously integrates pupillometry—an index of arousal and neuro-modulatory engagement (Joshi, Li et al. 2016, Wang, Baird et al. 2018, Joshi and Gold 2020, Joshi 2021)—with detailed behavioral, cognitive, and clinical assessments. As such, this approach offers a systems-level characterization of how attentional control mechanisms operate throughout childhood and adolescence. We further characterize whether these mechanisms differ for leftward versus rightward attention—a well-established asymmetry in behavior and brain organization whose electrophysiological basis remains unclear.

## Methods

### Participants

Data from 27 participants were initially collected. Of these, four participants were excluded prior to preprocessing due to eye-tracking equipment incompatibilities (e.g., calibration failure due to glasses).

Following EEG and eye-tracking preprocessing, one participant was excluded due to excessive EEG noise (>15% bad channels or >50% bad epochs), and an additional participant was excluded due to excessive saccades (>50% of trials exceeding 2° outside the central fixation region of interest; see *Eye-Tracking Preprocessing*). Subsequent analyses included a final sample size of 21, as described in **Table 1**.

**Table 1.** Participant demographics. Continuous variables: mean ± standard deviation. Categorical variables: n (%). Missing clinical data: IQ (n=19, missing 2); SRS-2 (n=18; missing 3); CPT-3 (n=20; missing 1) *IQ – intelligence quotient; SRS-2 – Social Responsiveness Scale, 2^nd^ edition; CPT-3 – Conners Continuous Performance Task, 3^rd^ edition*

|  | **TD (n=21)** |
| --- | --- |
| **Age, years** | 14.747 ± 3.776 |
| **Sex, male** | 10 (47.62%) |
| **Handedness** |  |
| Right | 19 (90.48%) |
| Left | 1 (4.76%) |
| Ambidextrous | 1 (4.76%) |
| **Full-scale IQ** | 106.579 ± 19.001 |
| **SRS-2** | 47.722 ± 7.622 |
| **CPT-3 Response Style** | 55.25 ± 11.571 |

Participants were recruited without regard to sex, race, or ethnicity; Intelligence quotients for verbal (V-IQ), and full-scale (FS-IQ) intelligence were assessed for all participants, using the Wechsler Abbreviated Scales of Intelligence (WASI-II; (Weschler 2011)) and Wechsler Intelligence Scale for Children (WISC-V; (Weschler 2014)).

The Social Responsiveness Score, 2^nd^ Edition (SRS-2; (Constantino, Davis et al. 2003, J.N. 2012)) questionnaire was collected from all participants to obtain a continuous measure of autistic traits. The SRS-2 yields age- and sex-normed T-scores (M = 50, SD = 10), with higher scores indicating higher levels of autistic traits. T-scores ≤59 are considered within the typical range, 60–65 indicates mild impairment, 66–75 moderate impairment, and ≥76 severe impairment consistent with clinically significant symptomatology.

Continuous measures of attention were obtained from Conners’ Continuous Performance Test, 3^rd^ edition (CPT-3; (Conners 2008)), which is a brief, computerized test to evaluate inattentiveness, impulsivity, sustained attention and vigilance in individuals >8 years old. As measured by the CPT-3, the Response Style Index is a signal detection statistic that reflects an individual’s natural response tendency in tasks involving a speed–accuracy trade-off. T-scores are age-normed (M = 50, SD = 10). A conservative response style (T ≥ 60) reflects an emphasis on accuracy over speed (i.e., a more cautious approach), whereas a liberal response style (T ≤ 40) reflects an emphasis on speed over accuracy (i.e., a more impulsive approach). T-scores between 41–59 indicate a balanced response style, reflecting sensitivity to both speed and accuracy.

To be included, participants had to have no history of neurological, developmental, or psychiatric disorders and be in age-appropriate grade at school when applicable. Exclusionary criteria included: (1) exclusionary physical limitations (e.g. significantly impaired and uncorrected vision), (2) were born prematurely (<35 weeks) and/or experienced prenatal/perinatal complications, or (3) FS-IQ <80.

All participants provided informed consent or assent in accordance with age-appropriate procedures approved by the Institutional Review Board of the Albert Einstein College of Medicine. For participants under 18 years of age, written parental or guardian consent was obtained in addition to participant assent. Participants 18 years or older provided their own written informed consent Participants received modest recompense for their participation (at $15 per hour).

### Procedure

Participants were seated in a double-walled, darkened, sound-attenuated, electrically shielded booth (International Acoustics Company, Bronx, NY) and gaze was stabilized using a chin rest. Visual stimuli were presented on an LCD monitor positioned 79 cm from the participant.

EEG recordings were collected from 64 active channels (10–20 system; at a digitization rate of 512 Hz, using Active Two amplifiers (BioSemi, Amsterdam, The Netherlands) with an anti-aliasing filter (−3 dB at 3.6 kHz). The recording reference was the Common Mode Sense (CMS) active electrode, which, together with the Driven Right Leg (DRL) passive electrode, forms a feedback loop to stabilize the reference signal and reduce common-mode noise. This referencing convention in BioSemi systems differs from traditional reference electrodes by dynamically compensating for potential differences between the scalp and the amplifier ground, effectively creating a "zero potential" point.

The paradigm was implemented in Python using PsychoPy, and analog triggers marking stimulus onset and button presses were transmitted to the acquisition computer via Lab Streaming Layer (LSL) (Kothe, Shirazi et al. 2024). Gaze position and pupil data were recorded using the EyeLink 1000 system (SR Research Ltd., Mississauga, Ontario, Canada) at a sampling rate of 500 Hz.

### Task

#### UDTR

All participants underwent a staircase procedure at the beginning of testing. This procedure, known as the Up-Down Transformed Rule (UDTR) was used to rapidly equate performance across the task and across participants (Wetherill and Levitt 1965) before the beginning of the formal experimental sessions. Participants were instructed to respond to the presence of a white target ring via keyboard press (left key = present; right key = absent). Correct responses triggered an adaptive staircase procedure that adjusted stimulus contrast based on accuracy. We used a 3-up, 1-down rule, meaning that, when a participant made three consecutive correct responses, the stimulus contrast level was decreased by one step, and for any incorrect response, stimulus contrast level was increased by one step. This set of rules converges on an accuracy level of 79.4%.

Importantly, although S2 stimuli were presented both unilaterally and bilaterally in the primary task, the UDTR procedure employed only right-sided, unisensory S2 stimuli, similar to the unisensory calibration procedure described by Murphy et al (Murphy, Foxe et al. 2016). As such, these thresholds do not account for the additional competitive demands introduced by a second, spatially distinct stimulus during the main task, or different thresholds based on hemifield.

#### Attention Task

We adapted a traditional spatial attention paradigm that has been previously applied in children of this age (Banerjee, Snyder et al. 2011), which consists of a simple Stimulus 1 (S1) – Stimulus 2 (S2) design. Each trial consisted of an instructional cue (S1), an intervening blank preparatory period (cue-S2 window), followed by a task-relevant second stimulus (S2).

The S1 is a centrally presented arrow (the cue; 1.4° in diameter) to indicate which side of space the subject should covertly attend, while remaining centrally fixated. The stimulus-onset-asynchrony (SOA) between the cue onset and S2 onset (i.e. the Cue–S2 period) was fixed at 1,120 ms. After the Cue to S2 interval, the S2s were presented for a duration of 160 ms. The cue remained in the center of the monitor until S2 offset. The visual S2 stimulus consisted of social (grayscale faces) and non-social (grayscale houses) images, subtending 9.15° [height] × 6.62° [width], centered 5.5° to the left and right of the fixation cross. House and face stimuli were matched in luminance, brightness, and contrast. All visual stimuli were presented on a gray background. The inclusion of both social and non-social stimuli allowed us to examine whether attentional sensory gain mechanisms differ as a function of stimulus social relevance.

The subject’s task was to monitor the stimulus on the attended (cued) side and to respond via keyboard-press when they detect a white target ring (inside diameter of 0.90°; outside diameter of 0.94°; combined by screen overlay on top of the house/face S2). While the target could appear on either, neither, or both sides, participants are instructed to respond only to targets on the attended side, ignoring task-irrelevant stimuli on the uncued side. During the experimental session, participants were instructed to respond as quickly and accurately as possible to targets on the cued side and to withhold responses otherwise. The likelihood of receiving a valid target (on the attended stimulus) was set at 33%. Target ring brightness was determined per-subject via the UDTR staircase method described above. Following S2, a 1,000 ms response period was presented. This was followed by a 1,000 ms fixation interval preceding the next cue, during which the fixation cross remained on the screen (see **Fig. 1a**). The task was performed in three contexts:

**Figure 1.**
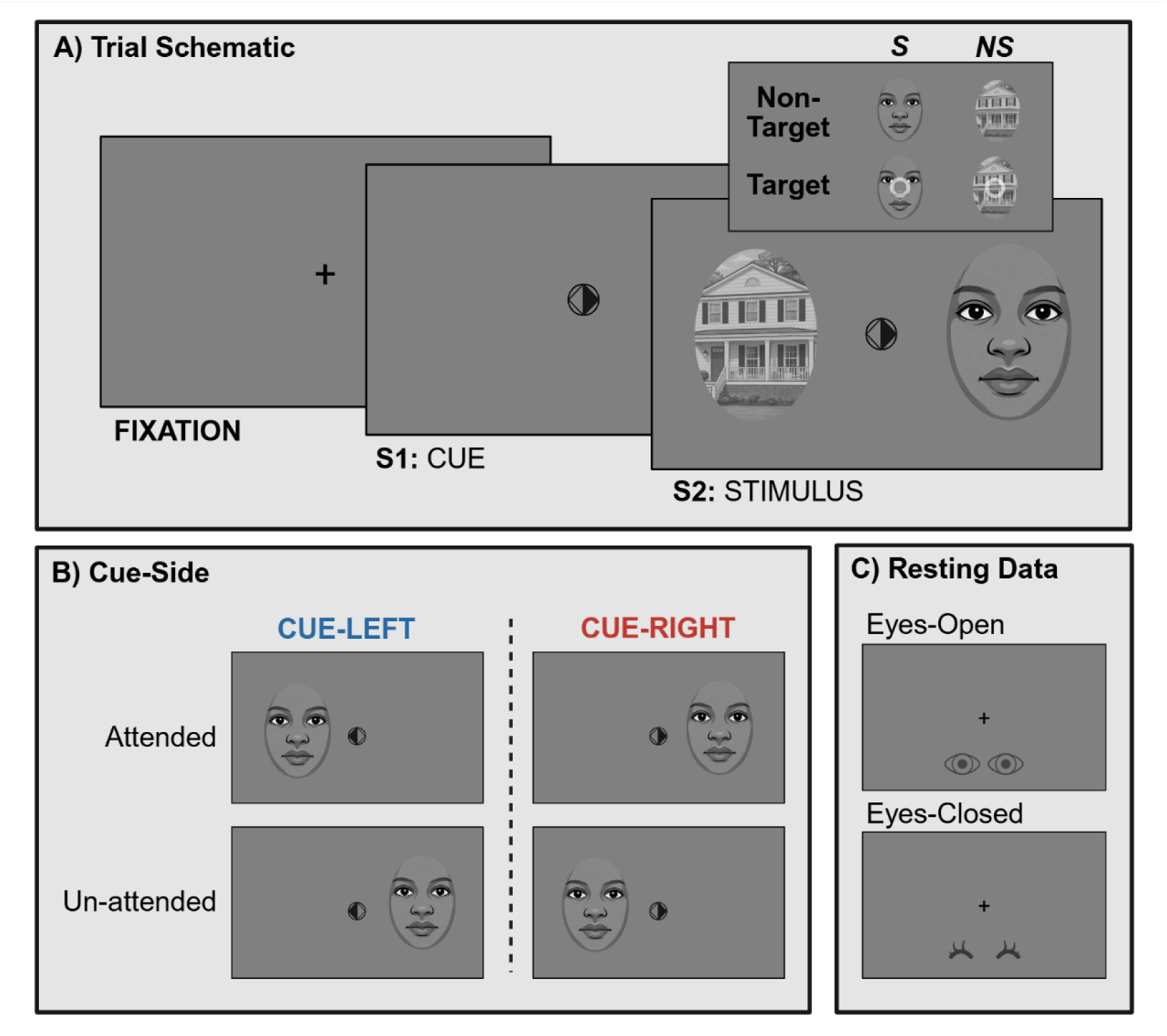
Experimental design. **a) Trial schematic.** Each trial began with a fixation period (1,000 ms), followed by a brief spatial cue (S1) indicating which side to attend to. After an anticipatory interval (1,120 ms), stimuli were presented (S2; 160 ms), consisting of social (S; faces) and/or non-social (NS; houses) images. Participants then responded via-keyboard press based on the presence of a white target ring on the attended side (response interval: 1,000 ms). This schematic is for illustrative purposes only and is not to scale. Face stimuli were originally real faces images (NimStim database); for display purposes here, we show stylized comic representations to avoid identifiable likenesses. **b) Cue-side.** Attention was directed to either the left (cue-left) or right (cue-right) visual hemifield via a central arrow cue. Attended refers to stimuli presented in the cued hemifield (left for cue-left; right for cue-right), whereas unattended stimuli appeared in the opposite hemifield. Only unilateral stimulus presentations are illustrated here; however, bilateral trials were also included, in which attended and unattended stimuli were presented simultaneously. **c) Resting data.** Resting-state EEG was collected both before and after the task to characterize participants’ baseline oscillatory activity. Each resting session consisted of 2 minutes of eyes-open and 2 minutes of eyes-closed recording.

**Context 1 (non-social):** S2s are **non-social** visual stimuli (house).

**Context 2 (social):** S2s are **social** visual stimuli (face).

**Context 3 (inter-mixed):** S2s are inter-mixed with **social** (faces) and **non-social** (houses) visual stimuli.

The first two conditions were blocked by social context (Context 1: non-social; Context 2: social), with order randomized across participants. In these “pure” contexts, S2 was presented under one of three conditions: unilaterally on the attended side, unilaterally on the unattended side, or bilaterally. Unilateral presentations occurred with a combined probability of 67% (attended and unattended equally likely), while bilateral presentations occurred with a probability of 33%. These blocks allow for evaluation of how top-down expectations of social and non-social stimuli influences attentional modulation.

Pure blocks were always followed by an inter-mixed condition (Context 3), where face and house S2 stimuli (attended and unattended) occur with equal probability. In inter-mixed blocks, S2 was presented under one of four conditions for both social and non-social stimuli, yielding a total of eight combinations: unilateral attended (S, NS), unilateral unattended (S, NS), bilateral with identical stimuli (S/S, NS/NS), or bilateral with different stimuli (S/NS, NS/S). Unilateral presentations occurred with a combined probability of 50% (attended and unattended equally likely), and bilateral presentations occurred with a probability of 50%. With social and non-social stimuli presented unpredictably within a single block, top-down contextual effects are not present and we were able to assess the potential contribution of bottom-up attentional biases induced by differences in stimulus type (i.e., social versus non-social stimuli). While this approach may introduce order effects, it prevented potential carry-over effects from the intermixed condition.

Each participant completed 40 blocks (9 pure social, 9 pure non-social, 22 inter-mixed) of 55 trials each, resulting in the collection of ∼167 trials per S2 combination. S2 combinations refer to the possible configurations at stimulus onset, including an attended stimulus presented alone, an attended stimulus presented with an unattended distractor, or an unattended distractor presented without an attended stimulus. Data collection for this paradigm took ∼4 hours with lunch and frequent short breaks, to maintain attention.

### Resting State

Resting-state EEG was recorded before and after task completion to characterize each participant’s intrinsic oscillatory profile (e.g., baseline alpha activity) and to assess how a period of effortful task engagement altered these dynamics. Each resting-state session consisted of two 2-minute blocks: an eyes-open block followed by an eyes-closed block. During the eyes-open block, participants viewed a uniform gray screen with a central fixation marker to support stable gaze and reduce movement-related artifacts. For the eyes-closed block, participants were instructed to close their eyes and remain still. The same resting-state protocol was repeated post-task. Because the difference between eyes-open and eyes-closed conditions offer a measure of alpha reactivity, both conditions were included in the analyses rather than selecting a single baseline state. Analyses specifically examining fatigue-related changes in oscillatory dynamics between pre- and post-task recordings will be interpreted in a separate study.

Three participants were excluded from the resting-state analysis due to poor data quality (>20% bad channels) in either the before or after block, yielding a final resting-state sample of *n* = 18.

### Pre-Processing

#### Eye-Tracking

Eye-tracking—which was exported using Data Viewer software Version 4.3.210—served two distinct purposes in the present study: (1) to confirm compliance with covert spatial attention instructions by ensuring central fixation was maintained, and (2) to measure task-evoked pupil responses as an index of arousal and cognitive effort.

##### Quality Control

Eye tracking data were cleaned individually for each participant using custom Python scripts and following best practices (Mathot 2018, Mathot and Vilotijevic 2023). Offscreen data were removed, blinks and artifacts were identified and corrected. After cleaning, trials missing over 50% of pupil data were excluded.

##### Removal of Saccade Trials

Additional pre-processing focused on gaze was completed to ensure stable central fixation at stimulus onset. To account for potential drift—which can introduce block-specific offsets in eye-tracking calibration—the center of fixation was estimated separately for each block. Specifically, for each block and participant, a baseline fixation window prior to cue onset (−1.22 to −1.12 s) was used to compute the median horizontal and vertical gaze position across all trials within that block. This block-specific median gaze position served as the estimated fixation center, allowing robust correction for gradual drift or posture-related calibration offsets.

For each trial, fixation stability was evaluated at stimulus onset (−100 to +100 ms). Gaze samples were classified as within fixation if their two-dimensional (x–y) distance from the block-specific fixation center fell within a circular region of interest (ROI) with a radius of approximately 2° of visual angle (∼2° radius). The ROI radius was selected based on stimulus geometry: face stimuli were centered 5.5° from fixation and subtended 6.62° in width, placing the inner edge of the stimulus at approximately 2.2° from fixation (5.5° − 3.31°). Thus, a 4° fixation window ensured that gaze remained within central fixation and did not encroach upon the nearest edge of the stimulus. The fixation center was defined as the median gaze position during the pre-cue baseline window. Trials were retained if at least 80% of valid (non-missing) gaze samples within this window fell inside the fixation ROI. Trials that failed to meet this criterion were excluded from further analysis.

At the participant level, individuals were excluded if fewer than 50% of trials met the fixation criterion, reflecting excessive saccadic behavior or unstable gaze that would compromise interpretation of attentional effects. This procedure ensured that retained trials reflected consistent central fixation required of a covert attention task.

#### EEG

EEG data were analyzed using Python (MNE; (Gramfort, Luessi et al. 2013)) and custom in-house scripts. Raw data were low-pass FIR-filtered at 40 Hz and high-pass FIR-filtered at 0.01 Hz.

Bad channels were detected automatically using *pyPrep.NoisyChannels* (Bigdely-Shamlo, Mullen et al. 2015), which includes a combination of classical bad-channel detecting functions: low signal-to-noise ratio, RANSAC (Martin A. Fischler 1981) prediction or correlation with other channels, abnormal amplitude or amounts of high-frequency noise, and near-flat channels. Subjects were excluded if >15% of channels were identified as bad; otherwise, bad channels were interpolated using the spherical-spline method (Perrin, Pernier et al. 1989).

Epochs were constructed from a −2,280 to +1,000 ms time window surrounding the onset of S2 (t=0 ms). Independent Component Analysis (ICA) using the *fastica* algorithm (max 1,000 iterations) was applied to epoched data high-pass filtered at 1 Hz to identify and manually remove components corresponding to eye movements, including horizontal saccades and blinks. The 1 Hz high-pass filter was applied only for ICA component identification, and all subsequent analyses were conducted on the original data (0.01-40 Hz) after applying ICA removal.

Epochs containing artifacts were excluded by automatic artifact rejection *(autoreject;* (Jas, Engemann et al. 2017)), which utilizes a cross-validation metric to identify an optimal amplitude threshold. Subjects were excluded if >50% of trials were rejected across all conditions.

Epochs were averaged at the individual subject level, for each of the stimulus conditions of interest, for subsequent analyses. Data was re-referenced to the average of all electrodes, a commonly used reference technique for evaluating event-related potentials (Tsuchimoto, Shibusawa et al. 2021).

### Analysis

#### Overview

Analyses were designed to characterize the neural mechanisms supporting anticipatory spatial attention and were organized around two temporal windows: (i) preparatory activity during the cue–S2 interval (−1000 to 0 ms), where anticipatory oscillatory dynamics were examined, and (ii) stimulus-evoked responses following S2 onset (0–1000 ms). Within the stimulus-evoked window, EEG and pupil responses were quantified to assess attentional modulation of early sensory gain. Within the preparatory interval, we examined oscillatory activity with a focus on parieto-occipital alpha-band dynamics associated with spatially selective attention. Finally, motivated by growing evidence that low-frequency frontal theta rhythms coordinate large-scale attentional control, we tested whether pre-stimulus theta phase predicts behavioral performance and modulates both posterior oscillatory activity and stimulus-evoked sensory responses.

#### ET: Evoked Response

##### Baseline Correction

Eye-tracking epochs were baseline-corrected using a pre-stimulus baseline window (−200 to −50 ms prior to S2 onset). Analyses focused exclusively on non-target trials to eliminate target- and motor-related confounds of the pupil response.

##### Extraction of Stimulus-Evoked Pupil Response

Peak amplitude of the stimulus-evoked pupil response was calculated as the maximum positive deflection within the S2 window (0 – 900 ms). Peak latency was extracted as the time point corresponding to this maximum value. Analysis of the pupil response were conducted only for unilateral stimulus configurations to assess attention effects without distractor confounds during bilateral stimulus presentation.

#### EEG: Evoked Response

##### Baseline Correction

Event-related potentials (ERPs) were analyzed from epoched EEG data time-locked to S2 onset. Epochs were extracted from −200 to 500 ms relative to stimulus onset and baseline-corrected using a pre-stimulus baseline window (−200 to −50 ms prior to S2 onset). Analyses focused exclusively on non-target trials to isolate attentional modulation of sensory-evoked activity whilst eliminating target and motor specific responses.

##### Extraction of ERP Components

In the present study, analyses of ERP components indexing attention focused specifically on the P1 component as an index of early sensory gain in extrastriate visual cortex (Awh, Anllo-Vento et al. 2000, Frey, Schmid et al. 2014).

Mean P1 amplitude was calculated as the average voltage within a 110–150 ms window centered on the grand-average P1 peak. Peak amplitude and latency were estimated within a broader 100–160 ms window, with peak amplitude defined as the maximum positive deflection and peak latency as the corresponding time point. ERP analyses were conducted separately for unilateral and bilateral stimulus configurations.

The P1 was extracted from bilateral parieto-occipital electrodes defined a priori based on canonical visual ERP topographies, and in line with the literature (Luck, Heinze et al. 1990, Heinze, Mangun et al. 1994, Wijers, Lange et al. 1997, Woldorff, Fox et al. 1997). Left occipital/parieto-occipital electrodes comprised O1, PO3, and PO7, and right occipital/parieto-occipital electrodes comprised O2, PO4, and PO8. Because the P1 indexes early sensory gain in extrastriate cortex, analyses focused on posterior occipital channels where the component is maximally expressed. For unilateral trials, P1 responses were quantified using a contralateral framework relative to stimulus location to capture maximal visual responses. Stimuli appearing in the right visual field were analyzed using left occipital electrodes (O1, PO3, PO7), whereas left visual field stimuli were analyzed using right occipital electrodes (O2, PO4, PO8).

#### EEG: Time-Frequency Analysis of Oscillatory Activity

Single-trial oscillatory power and phase were derived from complex time–frequency representations computed using complex Morlet wavelets (MNE-Python).

##### Anticipatory Alpha Power Extraction

To quantify anticipatory alpha activity, Morlet-based time–frequency power estimates were extracted in the alpha band (7–13-Hz) for each trial, channel, and time point. Alpha power was baseline-normalized using an MNE-style log-ratio transformation:

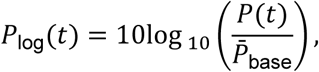

where *P̅*_base_ denotes the per-trial mean power within a pre-stimulus baseline window (−200 to −50 ms relative to cue onset).

Attentional modulation of alpha power is typically quantified within the cue–S2 anticipatory interval. In the present study, this window was defined as −800 to −200 ms prior to S2 onset, based on spatiotemporal cluster permutation results demonstrating maximal cue-related alpha modulation during this period (see *Spatiotemporal Cluster Permutation: Attentional Modulation of Anticipatory Alpha Power*; Supplementary Figure 1A). This window therefore reflects the empirically strongest phase of preparatory modulation. Alpha power was averaged across time within this window.

Anticipatory alpha power was extracted from parieto-occipital (PO) electrodes, defined a priori based on bilateral channel groups commonly used to index spatial attention–related alpha modulation. Specifically, the left PO ROI comprised *P3, P5, P7, PO3, PO7, O1*, and the right PO ROI comprised *P4, P6, P8, PO4, PO8, O2*. These ROIs were further corroborated by the spatial distribution of effects observed in independent sensor-level cluster-based permutation analyses (see *Spatiotemporal Cluster Permutation*, *Results*, and **Supplementary Figure 1A**).

To quantify spatially selective alpha modulation, PO activity was parameterized in a contralateral/ipsilateral framework. Trials were split by cue side (cue-left vs cue-right). For cue-left trials, the contralateral ROI (corresponding to the to-be-attended region) was defined as the right PO electrodes and the ipsilateral ROI (corresponding to the to-be-ignored, or unattended, region) as the left PO electrodes; for cue-right trials, this mapping was reversed.

##### Resting-State Spectral Analysis

To determine whether task-related changes in alpha were attributable to baseline spectral differences, resting-state alpha power was extracted from the same bilateral parieto-occipital ROIs described above.

To avoid edge artifacts introduced by time–frequency convolution, the first and last 10 seconds of each 120-second resting block were excluded, and analyses were restricted to the central 100-second interval. Morlet-based time–frequency decomposition (7–13-Hz) was applied identically to the task data. Alpha power was computed at the single-channel level and averaged within each hemisphere-specific PO ROI (left: *P3, P5, P7, PO3, PO7, O1*; right: *P4, P6, P8, PO4, PO8, O2*). Resting alpha power was quantified separately for hemisphere (left vs right), recording period (pre-task vs post-task), and eye condition (eyes-open vs eyes-closed), allowing direct comparison of baseline hemispheric spectral profiles independent of spatial attention demands.

In addition to Morlet-based power estimates, resting spectra were parameterized using the *specparam* (Donoghue, Haller et al. 2020) algorithm to dissociate periodic and aperiodic components of the power spectrum. For each ROI and condition, spectral fits yielded estimates of the aperiodic exponent and alpha-band peak center frequency. This parameterization allowed assessment of whether any hemispheric differences were attributable to oscillatory peak characteristics or to shifts in the underlying aperiodic background activity.

#### EEG: Phase-Behavior Relationship

To explore whether intrinsic oscillatory phase predicts behavioral performance, we examined, on a trial-by-trial basis, if the phase of ongoing oscillatory activity during the cue-S2 interval (when ongoing oscillatory activity reflects preparatory attentional state prior to stimulus onset) was associated with the RT to the S2 target. Given the exploratory and hypothesis-generating nature of these analyses, phase–behavior relationships were assessed across all scalp channels and frequencies below 15 Hz without a priori spatial or spectral restriction.

Statistical significance was evaluated using nonparametric permutation-based p-values without multiple testing correction. To guard against false positives and control family-wise error, inference relied on cluster-based permutation procedures, which assess effects at the level of spatially and/or spectrally contiguous clusters rather than individual comparisons. This inherently conservative framework substantially reduces the likelihood of spurious findings arising from isolated sensors or frequencies (Maris and Oostenveld 2007). Still, all results are interpreted cautiously and are intended to inform subsequent targeted analyses.

##### Instantaneous Phase Extraction

To obtain frequency-resolved phase estimates, we assessed frequencies spanning 3–15 Hz. For each frequency ***f***_***c***_, we selected the corresponding wavelet frequency bin from the complex time–frequency coefficients. Phase at each frequency bin was computed as the angle of the complex signal. This yielded instantaneous phase time series for each trial and channel within the epoch.

##### Phase and RT Binning

All analyses were conducted separately for cue-left and cue-right trials. Analyses were limited to attended target trials, and only hit trials were included when computing phase–RT functions, such that phase–behavior coupling reflects modulation of RT magnitude among correct responses. Trials with missing responses were excluded for subsequent analyses.

For each channel and cue condition, phase–behavior relationships were computed by binning trials according to pre-stimulus phase using a sliding circular window. Pre-stimulus phase was sampled at a frequency-specific time point defined by the Morlet wavelet length, specifically the wavelet center time such that the end of the wavelet aligned with stimulus onset (t = 0 s), consistent with the phase–behavior framework described by Fiebelkorn et al (Fiebelkorn, Pinsk et al. 2018). This separation minimizes contamination from cue- or S2-evoked responses while isolating ongoing preparatory oscillatory activity. For each frequency we computed:

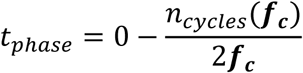

where 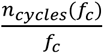 reflects the temporal duration of the Morlet wavelet, and *t_Phase_* corresponds to the center of a wavelet whose final edge aligns with stimulus onset (t = 0).

Phase bins were centered every 10° across the full 360° cycle, with each bin spanning a 120° window. For each bin, mean RT (RT) was calculated among all hit trials falling within that phase window. Bins without sufficient data (<20 trials) were set to 0% change, indicating no deviation from baseline performance.

To normalize for baseline performance differences, RT was expressed as percent change relative to the overall RT for that cue condition:

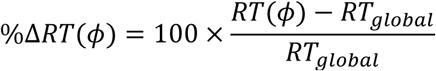

where RT(ϕ) reflects mean RT among hit trials in phase bin ϕ, and RT_global_ reflects the global mean RT across all included hit trials for that channel and cue condition.

##### Quantifying the Phase-RT Relationship: Fast Fourier Transform

For each channel and cue condition, rhythmic modulation of RT was tested by extracting the amplitude of the fundamental (1-cycle) component of the phase–behavior function.

To account for physiologically plausible volume conduction that may spatially blur phase–behavior relationships across neighboring electrodes, spatial smoothing was applied to the observed data by conservatively averaging the phase-specific hit rates at each electrode with those of its four closest neighboring electrodes, following the approach described by Fiebelkorn et al. (Fiebelkorn, Snyder et al. 2013). For each channel, smoothed curves were computed by averaging the percent-change RT functions across the channel itself and its four nearest neighboring electrodes, defined based on Euclidean distance in 2-D sensor space derived from the EEG montage.

Smoothed percent-change curves were demeaned and transformed using the discrete Fourier transform. The magnitude of the first harmonic was converted to amplitude in percent-change RT units using standard FFT normalization:

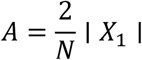

where *X*_1_ is the first non-zero Fourier coefficient and *N* is the number of phase bins. This amplitude reflects the strength of sinusoidal modulation of behavioral performance across the phase cycle.

##### Permutation Testing

Statistical significance of phase–behavior coupling was assessed using a nonparametric permutation procedure. For each channel, cue condition, and frequency, behavioral outcomes (RT) were randomly shuffled relative to phase values for 1,500 permutations, as recommended by Pernet et al. (Pernet, Latinus et al. 2015). For each permutation, phase–hit-rate functions were recomputed and the 1-cycle FFT amplitude was extracted using the same procedure as for the observed data.

P-values were computed as the proportion of permutation amplitudes greater than or equal to the observed amplitude:

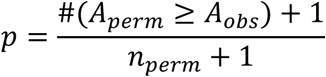

Statistical significance of channels exhibiting a significant phase-behavior relationship was assessed at p < 0.05 without correction for multiple comparisons across channels, consistent with an exploratory sensor-level analysis.

#### EEG: Phase-Power Relationship

We next examined whether pre-stimulus theta phase modulates parieto-occipital alpha (7–13-Hz) and beta (13–20 Hz) oscillatory power in a behaviorally relevant manner, restricting analyses to frequencies and channels that exhibited significant phase–behavior coupling in the preceding analysis.

We first assessed phase-dependent modulation of both alpha and beta power, quantifying effects separately for parieto-occipital regions corresponding to the attended versus unattended space. This allowed us to determine whether theta phase modulated bilateral or unilateral posterior oscillatory activity under cue-left and cue-right conditions.

Phase–power relationships were computed separately for cue-left and cue-right trials using the identical sliding-window phase-binning procedure described above. Trials were binned according to pre-stimulus theta phase using overlapping 120° windows stepped every 10° across the full 360° cycle. Importantly, in contrast to the phase–RT analysis, all valid trials (including non-targets) were included regardless of behavioral outcome, as the goal was to maximize statistical power to detect preparatory cross-frequency interactions independent of performance variability. For each bin, mean power was calculated across all trials falling within the phase window.

Power modulation was expressed as percent change relative to the global mean power for the corresponding cue condition and frequency:

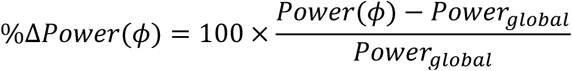

Phase-binned power curves were demeaned and transformed using the discrete Fourier transform, and the amplitude of the first harmonic was extracted using the same normalization applied in the phase–behavior analysis. To test whether phase-dependent fluctuations in power were statistically significant, we used the same cluster permutation procedure described above (see *Permutation Testing*).

To determine whether the effect was attributable to broadband alpha modulation, or instead was driven by a narrower, frequency-specific component within the alpha band, we further examined whether phase-dependent modulation was spectrally focal within the alpha range by repeating the analysis at each individual frequency (7–13-Hz), using the identical phase-binning and FFT-based quantification procedure.

#### EEG: Phase-P1 Relationship

To determine whether pre-stimulus theta phase modulates early sensory-evoked responses, we quantified the relationship between phase and peak P1 amplitude using the same phase-resolved framework described above. Analyses were conducted separately for cue-left and cue-right conditions, and for parieto-occipital regions corresponding to the attended versus unattended space.

Trials were sorted by pre-stimulus theta phase using overlapping 120° windows stepped every 10° across the full 360° cycle. Within each phase window, P1 amplitudes were averaged across trials to generate phase-binned response functions.

Modulation of P1 amplitude was expressed as percent change relative to the condition-specific global mean:

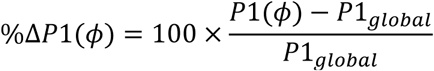

The resulting phase-binned curves were demeaned and analyzed in the frequency domain using the discrete Fourier transform, and the amplitude of the first harmonic was extracted using the same normalization as in the phase–behavior and phase–power analyses. Statistical significance of phase-dependent P1 fluctuations was evaluated using the identical permutation-based testing procedure described above (see *Permutation Testing*).

### Statistical Testing

#### Per-Participant Linear Mixed-Effects Modeling: Behavioral and Neural Effects

Statistical analysis of behavioral and neural measures per-participant was conducted using linear mixed-effects modeling (LMM; (Bates D 2015)) in R version 4.4.0 using the *lme4* package. Follow-up pairwise comparisons were obtained by estimated marginal means (*emmeans*; (Lenth 2019)) to compute contrasts, applying a Tukey correction. All primary outcome measures were analyzed using LMMs with subject-specific random intercepts and Age as a covariate; fixed-effect structures for each analysis are detailed in **Table 2**.

**Table 2.** LMM specifications for behavioral, pupillometric, neural, and spectral analyses. The table summarizes dependent variables and corresponding fixed-effect predictors for each primary analysis. All models included age as a covariate and subject as a random intercept (1 | Subject).

| Measure | Predictors | Results |
| --- | --- | --- |
| **Behavior** (RT, hit rate, false alarm rate, $d'$, IES) | S2 Type (unilateral/bilateral) × Block (S/NS/IM) × Cue Side (L/R) + Age + (1 \| Subject) | Table 3, Fig. 2 |
| **Stimulus-evoked pupil response** | Attention (attended/unattended) × Condition (S/NS) × Cue Side (L/R) + Age + (1 \| Subject) | Table 4, Fig. 3 |
| **Stimulus-evoked neural response** | Attention (attended/unattended) × Condition (S/NS) × Cue Side (L/R) + Age + (1 \| Subject) | Table 5, Fig. 4 |
| **Anticipatory neuro-oscillatory power** (alpha, beta) | Attention (unattended/attended ROI) × Block (S/NS/IM) × Cue Side (L/R) + Age + (1 \| Subject) | Table 6, Fig. 5 |
| **Resting alpha** (power, frequency, aperiodic exponent) | Hemisphere (left/right parieto-occipital ROI) × Condition (eyes-open/eyes-closed) × Resting-State (RS)-Block (before-task/after-task) + Age + (1 \| Subject) | Table 7, Fig. 6 |

#### Single-Trial Linear Mixed-Effects Modeling: Alpha–P1 Contributions to RT

To determine whether pre-stimulus alpha power and early sensory gain independently predicted behavioral performance, we fit single-trial linear mixed-effects regression models with RT as the dependent variable. Trial-wise alpha power and P1 amplitude were entered simultaneously as fixed effects, with subject included as a random intercept to account for repeated measures. Alpha and P1 amplitude were z-scored within each group and cue direction prior to modeling to facilitate comparison of standardized effect sizes.

These models were implemented in Python (*statsmodels* v0.14; *mixedlm* function). Separate models were estimated for cue direction (left, right), as well as for hemispheric analyses of the attended ROI (target-side) and unattended ROI (distractor-side). This modeling framework allowed us to test whether oscillatory state (alpha) and early sensory gain (P1) explained unique variance in RT, and whether their behavioral contributions differed as a function of attentional direction and social context.

#### Spatiotemporal Cluster Permutation

In addition to the temporally and spatially constrained analyses above, spatiotemporal cluster permutation testing—in which all timepoints and channels were considered—was performed to provide a more comprehensive consideration of potential selective attention effects on both alpha power (**Supplementary Fig. 1A**) and the stimulus-evoked response (**Supplementary Fig. 1B**).

For cluster permutation testing of the evoked response, subject-level difference waves (attended − unattended) were entered into a one-sample spatiotemporal cluster permutation test against zero using MNE’s implementation. Attended trials correspond to stimuli presented in the cued hemifield, whereas unattended trials correspond to stimuli in the uncued hemifield.

To test attentional modulation of alpha power during the anticipatory window, subject-level cue-left minus cue-right difference maps (channels × time) were entered into a one-sample spatiotemporal cluster permutation test against zero, also using MNE’s implementation.

For both tests, clusters were defined as contiguous samples across both sensors and time exceeding a cluster-forming threshold corresponding to α = 0.05 (two-sided) at the sample level. Cluster-level significance was assessed using 1,500 permutations, with p-values computed based on the maximum cluster statistic to control the family-wise error rate across all sensors and time points. This approach provides stringent control against false positives by evaluating effects at the level of spatially and temporally coherent patterns rather than isolated sensors or time samples. The resulting cluster maps provided convergent support for the a-priori selection of occipital/parieto-occipital channels and component-specific time windows used in ROI-based ERP extraction.

#### Age-Related Changes in Alpha Asymmetry

Associations between neural measures and age were assessed using linear regression analyses. Alpha modulation was defined as the difference in log-transformed alpha power between the unattended and attended regions of interest (ROI), such that higher values indicate greater alpha power over the unattended relative to the attended ROI. Cue-side asymmetry in alpha modulation was calculated as the difference between leftward and rightward alpha modulation values (left − right). Thus, larger asymmetry values indicate stronger attentional modulation of alpha power during leftward compared with rightward attention.

Alpha asymmetry values were then regressed against age using ordinary least squares regression. Scatterplots display fitted regression lines with 95% confidence intervals, and regression coefficients (β), p-values, and model R² are reported.

## Results

### Behavior

As described in **Table 3**, LMM analyses revealed a significant main effect of cue side on RT (*F* = 3.96, *p* = 0.048). Follow-up testing indicated faster responses for Cue Left relative to Cue Right trials (Cue Left − Cue Right: *B* = −0.014, *p* = 0.031), as shown in **Fig. 2i**. Older participants also responded faster (*B =* −0.11, p=0.035). No significant main effects of S2 type (unilateral vs. bilateral), or block type (S, NS, IM) were observed for RT.

**Figure 2.**
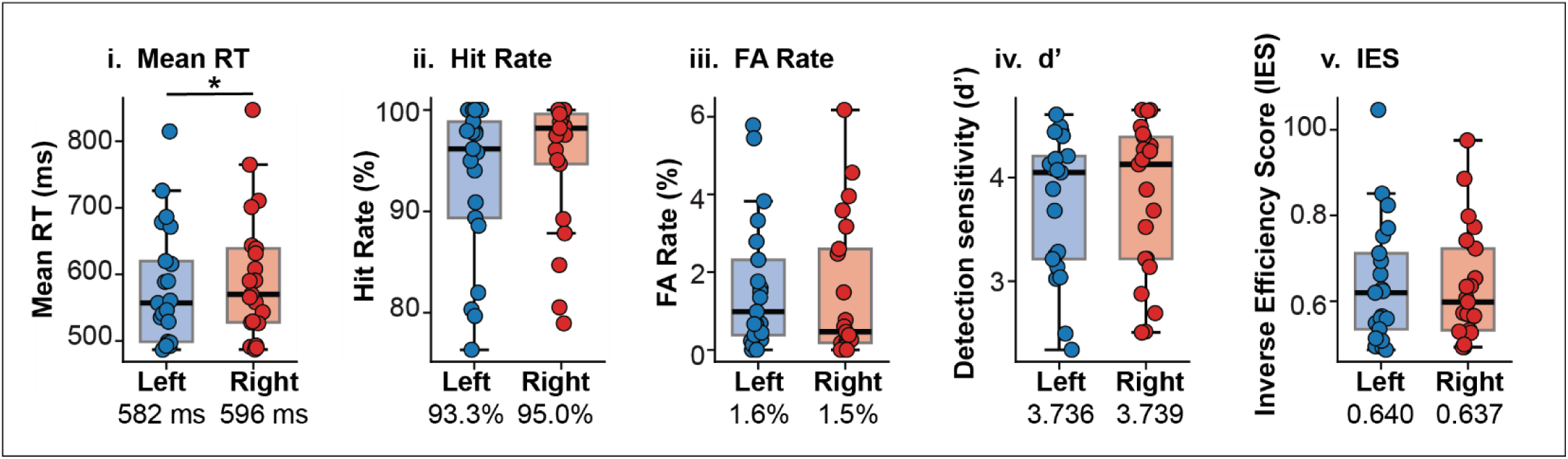
Cue-side effects on behavior, demonstrating faster RTs (RT) during leftward compared to rightward attention. Per-participant task performance is shown as **(i)** mean RT (RT), **(ii)** hit rate, **(iii)** false alarm (FA) rate, **(iv)** detection sensitivity (d’) and **(v)** inverse efficiency score (IES), plotted separately for Cue Left (blue) and Cue Right (red) trials, collapsed across stimulus category and block type. Each dot represents RT for a single participant; boxplots indicate the group distribution (median and interquartile range). The group mean for each measure and cue side is displayed beneath each boxplot. Asterisks denote a significant main effect of cue side (left vs. right) in behavioral performance (p<0.05).

**Table 3:** Behavioral performance. Summary of LMM results for behavioral performance, including measures of reaction-time (RT), hit rate, inverse efficiency score (IES), false alarm rate, and detection sensitivity (d’). For each participant, mean RT (RT) was calculated from correct target trials (hits only). Proportion of hits and misses were defined as the presence or absence of a response on target trials, respectively. False-alarm rate was computed as the proportion of non-target trials with a response. IES was calculated as mean RT divided by hit rate. d′ was computed using log-linear–corrected hit and false-alarm rates to avoid ceiling and floor effects. All measures were computed at the participant level. LMMs included fixed effects of S2 Type (unilateral/bilateral) × Block (S/NS/IM) × Cue Side (L/R), with Age entered as a covariate, and a random intercept for Subject *(1 | Subject)*.

|  | Cue Side Effects | S2 Type Effects | Block Effects | Age Effects |
| --- | --- | --- | --- | --- |
| RT | Main effect of Cue Side; driven by faster RT when attention is directed to the left | None observed | None observed | Main effect of Age; driven by faster RT in older subjects |
| Hits | None observed | None observed | None observed | Main effect of Age; driven by higher hit rate in older subjects |
| IES | None observed | None observed | None observed | Main effect of Age; driven by lower IES in older subjects |
| False Alarms | None observed | Main effect of S2 Type; driven by higher false alarm rate to bilateral (vs. unilateral) stimuli | None observed | Main effect of Age; driven by lower false alarm rate in older subjects |
| d' | None observed | None observed | None observed | Main effect of Age; driven by higher detection sensitivity in older subjects |

Hit rate and inverse efficiency score (IES) did not show significant effects of cue side, S2 type, or block type; however, both measures exhibited significant age-related effects, with increasing age associated with higher hit rates (*B* = 0.01, *p* = 0.013) and lower IES (*B* = −0.021, *p* = 0.006).

False-alarm rate showed a main effect of S2 type (*F* = 56.18, *p* < 0.001), indicating decreased false alarms in the presence of unilateral (vs. bilateral) S2 presentation (unilateral – bilateral: *B* = −0.017, p<0.001), as well as a significant effect of age (*B* = −0.017, *p* < 0.001), with fewer false alarms at older ages. Older participants also demonstrated higher detection sensitivity (*B* = 0.124, *p* < 0.001).

### S2-Evoked Pupil Response

Pupil responses showed no significant main effects of attention or social context in either the pure or intermixed conditions (**Table 4**). In contrast, a significant main effect of cue side was observed in both the pure (F = 7.937, p = 0.006) and intermixed (F = 6.783, p = 0.01) conditions (**Fig. 3**). Follow-up analyses indicated that this effect was driven by larger pupil size when attention was directed to the right compared with the left (left_pure_ – right_pure_: *B* = −9.677, p = 0.006; left_inter-mixed_ – right_inter-mixed_: *B* = −8.238, p = 0.01). No significant age effects were observed.

**Figure 3.**
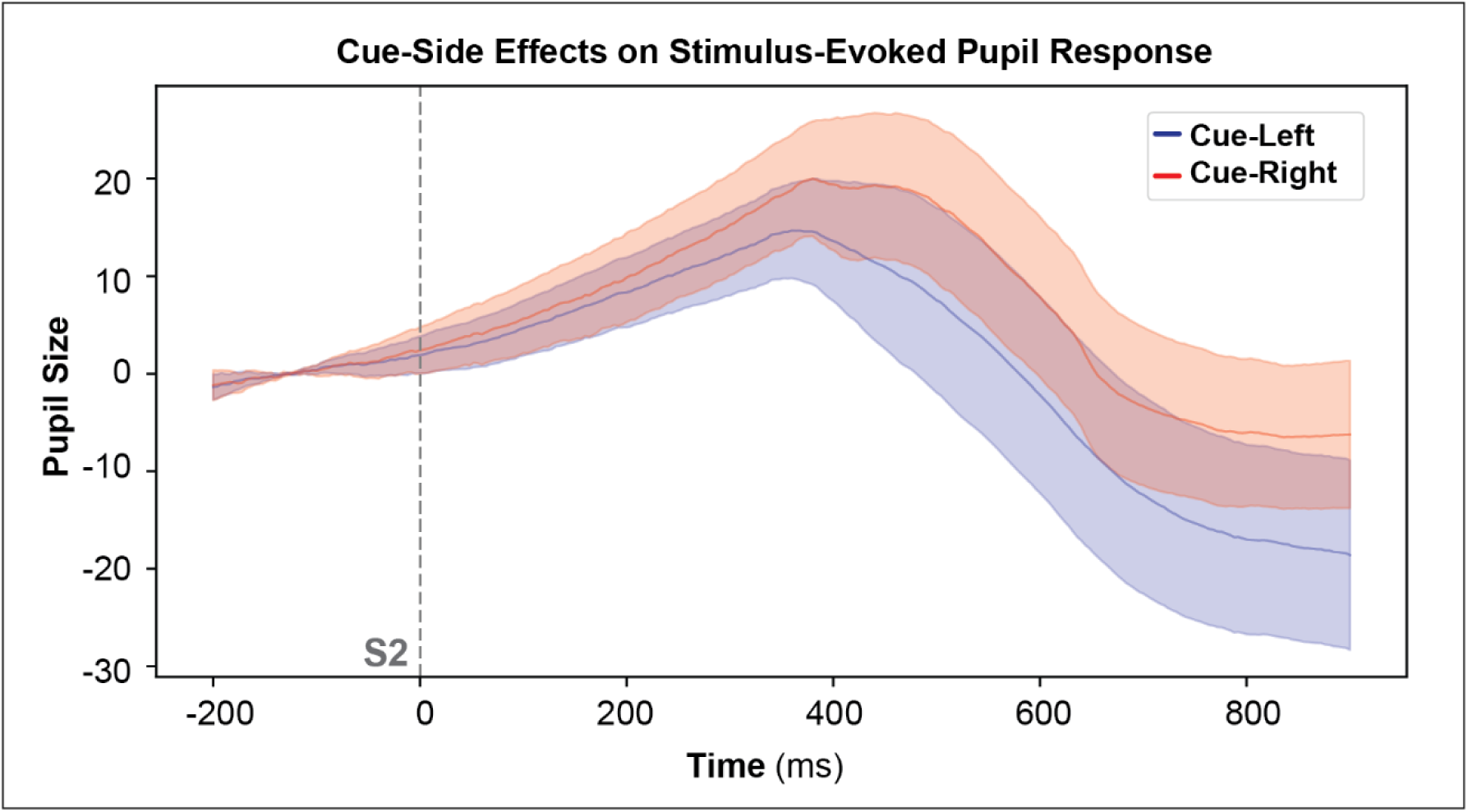
Larger stimulus-evoked pupil response during rightward attention. Grand-average stimulus-evoked pupil responses to unilateral non-target stimuli are shown for Cue Left (blue) and Cue Right (red) trials, collapsed across pure and intermixed blocks. The solid lines represent the group mean, with shaded regions indicating ±SEM. Time 0 (dashed vertical line) denotes S2 onset. All unilateral stimulus presentations are collapsed across attentional status, as no significant differences in pupil response were observed between attended and unattended stimuli.

**Table 4:** Pupil response to unilateral stimuli. Summary of LMM results for S2-evoked pupil response in the pure (S/NS) condition for non-target unilateral stimuli. Analyses were restricted to non-target trials to eliminate motor-response–related confounds. LMM included fixed effects of Attention (attended/unattended) × Condition (S/NS) × Cue Side (L/R), with Age entered as a covariate, and a random intercept for Subject *(1 | Subject)*.

|  |  | Attention Effects | Social Context Effects | Cue Side Effects | Age Effects |
| --- | --- | --- | --- | --- | --- |
| Pupil | Pure | None observed | None observed | Main effect of Cue Side; driven by larger pupil response when attention is directed to the right (vs. left) | None observed |
|  | Inter-mixed |  |  |  |  |

### S2-Evoked Neural Response

Evaluation of the S2-evoked neural response was conducted using LMMs. The results reported here (see **Table 5**) pertain to non-target unilateral stimuli. Analyses evaluating the evoked response to bilateral stimuli are not reported here, as those models were designed to specifically evaluate social/non-social attentional effects of distractor stimuli and are beyond the scope of the present manuscript; these analyses will be presented in a future report.

**Table 5:** Evoked response to unilateral stimuli. Summary of LMM results for S2-evoked P1 component for non-target unilateral stimuli. Analyses were restricted to non-target trials to eliminate motor-response–related confounds. LMM included fixed effects of Attention (attended/unattended) × Condition (S/NS) × Cue Side (L/R), with Age entered as a covariate, and a random intercept for Subject *(1 | Subject)*.

|  |  | Attention |  | Social Context Effects | Age Effects |
| --- | --- | --- | --- | --- | --- |
|  |  | Attention Effects | Attention x Cue-Side Interaction |  |  |
| P1 | Amplitude | Main effect of Attention; driven by greater (positive-going) amplitude to attended stimuli | *None observed* | *None observed* | Main effect of Age; driven by lower (positive-going) amplitude in older subjects |
|  | Latency | *None observed* | Attention x cue-side; driven by earlier latency in response to attended stimuli in left-directed vs. right-directed attention | Main effect of Condition; driven by earlier latency in response to social stimuli | Main effect of Age; driven by earlier latency responses in older subjects |

A robust main effect sign of attention was observed for the P1 component of the evoked response, for both mean (*F* = 19.149, *p* < 0.001; **Fig. 4c**) and peak (*F* = 20.492, *p* < 0.001) amplitude. Post hoc analyses indicated that these effects were driven by significantly greater amplitudes for stimuli presented to attended versus unattended locations for both mean (attend – unattend: *B* = 1.617, p < 0.001) and peak (attend – unattend: *B* = 1.646, p < 0.001) amplitude measurements.

**Figure 4.**
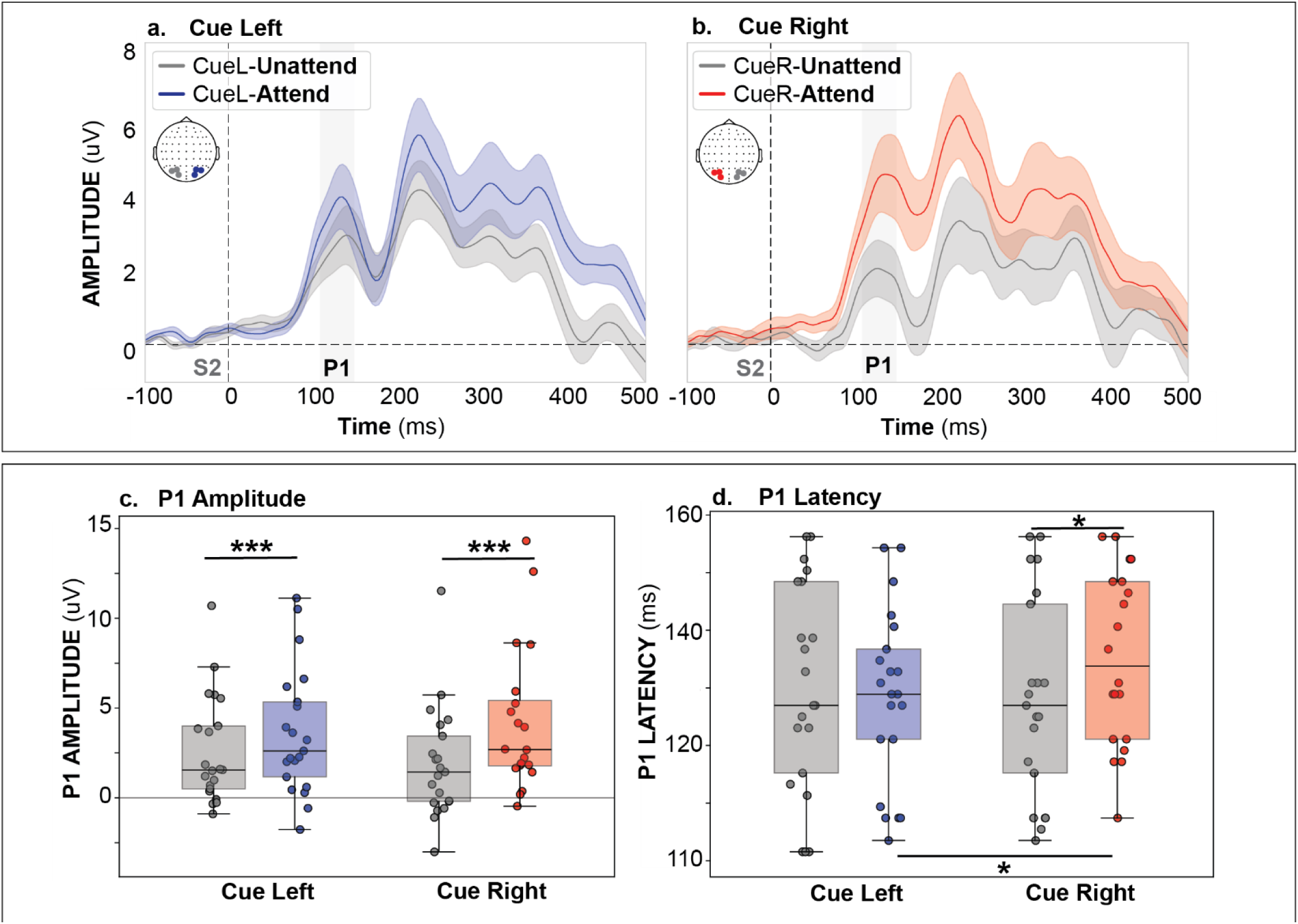
Cue-side effects on attentional modulation of the stimulus-evoked response. **a-b.** Grand-average event-related potentials (ERPs) elicited by unilateral non-targets are shown for Cue Left **(a)** and Cue Right **(b)** trials, collapsed across pure and inter-mixed blocks. Attended (blue: cue-left; red: cue-right) and unattended (gray) waveforms are overlaid, with shaded regions representing ±SEM. Attended responses correspond to stimuli presented in the cued hemifield, whereas unattended responses correspond to stimuli in the uncued hemifield. Time 0 denotes S2 onset (dashed vertical line). The P1 component, indexing early sensory attentional gain, is highlighted by the shaded gray time window (110–150 ms). **c-d.** Boxplots beneath each ERP panel summarize mean P1 amplitude (left) and latency (right) for attended (blue: cue-left; red: cue-right) and unattended (gray) unilateral non-targets. Asterisks indicate significant attention (attended vs. unattended) or cue side (left vs. right) (*p < 0.05, **p < 0.01, ***p < 0.001).

A significant attention × cue-side interaction was observed for P1 latency (*F* = 4.454, *p* = 0.037), indicating lateralized attentional modulation. Post hoc testing (**Fig. 4d**) revealed that the effect was driven by shorter latencies to attended stimuli presented on the left versus the right (attended: left – right. *B* = −0.007, *p* – 0.021). Only in rightward, but not leftward, attention was there a significant effect of attention on P1 latency (rightward: attend – unattend: *B* = 0.008, p = 0.014), which indicated a shorter latency in response to unattended stimuli (presented on the left) versus attended stimuli (presented on the right). No attention × cue-side interactions were observed for either P1 peak or mean amplitude.

A significant main effect of social context was also observed for P1 latency (*F* = 23.748, *p* < 0.001), driven by earlier P1 latencies to social compared with non-social stimuli (social – non-social: *B* = −0.011, p < 0.001).

Finally, age was significantly associated with mean amplitude (*B* = −0.485, p = 0.001), peak amplitude (*B* = - 0.581, p < 0.001) and latency (*B* = −0.002, p = 0.004) of the P1 component, revealing earlier latencies and lower amplitude P1 responses in older participants.

### Anticipatory Neuro-oscillatory Power

As shown in **Table 6**, analyses of neuro-oscillatory activity in the anticipatory window revealed a significant main effect of Attention for both alpha (F = 14.91, p < 0.001) and beta (F = 10.129, p = 0.002) power, in which power was higher over unattended versus attended ROI (alpha: *B* = 0.272, p < 0.001; beta: *B* = 0.162, p = 0.002).

**Table 6:** Anticipatory neuro-oscillatory activity. Summary of LMM results for anticipatory alpha and beta power (−800 to −200 ms) for all trials. LMM included fixed effects of Attention (unattended/attended ROI) × Block (S/NS/IM) × Cue Side (L/R), with Age entered as a covariate, and a random intercept for Subject *(1 | Subject)*.

|  | Attention |  | Block Effects | Age Effects |
| --- | --- | --- | --- | --- |
|  | Attention Effects | Attention: Interaction between Attention and Cue Side |  |  |
| Alpha power | Main effect of Attention; driven by greater alpha power over unattended ROI (vs. attended ROI) | Attention × Cue Side; driven by greater alpha power over unattended ROI (vs. attended ROI), only when attention is directed to the left | Main effect of Block; driven by lower alpha power in the inter-mixed blocks | None observed |
| Beta power |  |  |  | Main effect of Age; driven by lower beta power in older subjects |

This effect was qualified by a significant Attention × Cue Side interaction for both alpha (F = 13.945, p < 0.001) and beta (F = 16.23, p < 0.001) bands.

Follow-up analyses of alpha-power indicated that the attention effect was driven by greater power over the unattended versus attended ROI (**Fig. 5d**), but only when attention was directed to the left visual field (left: unattended – attended. β = 0.535, p < 0.001). Significant cue-direction differences were observed over both regions of interest, albeit in opposite directions. Alpha power over the unattended ROI was higher for leftward (versus rightward) attention (unattended: left – right. β = 0.244, p = 0.015), while alpha power over the attended ROI was lower in leftward (versus rightward) attention (attended: left – right. β = −0.282, p = 0.005), indicating bilateral modulation over both unattended and attended ROI.

**Figure 5.**
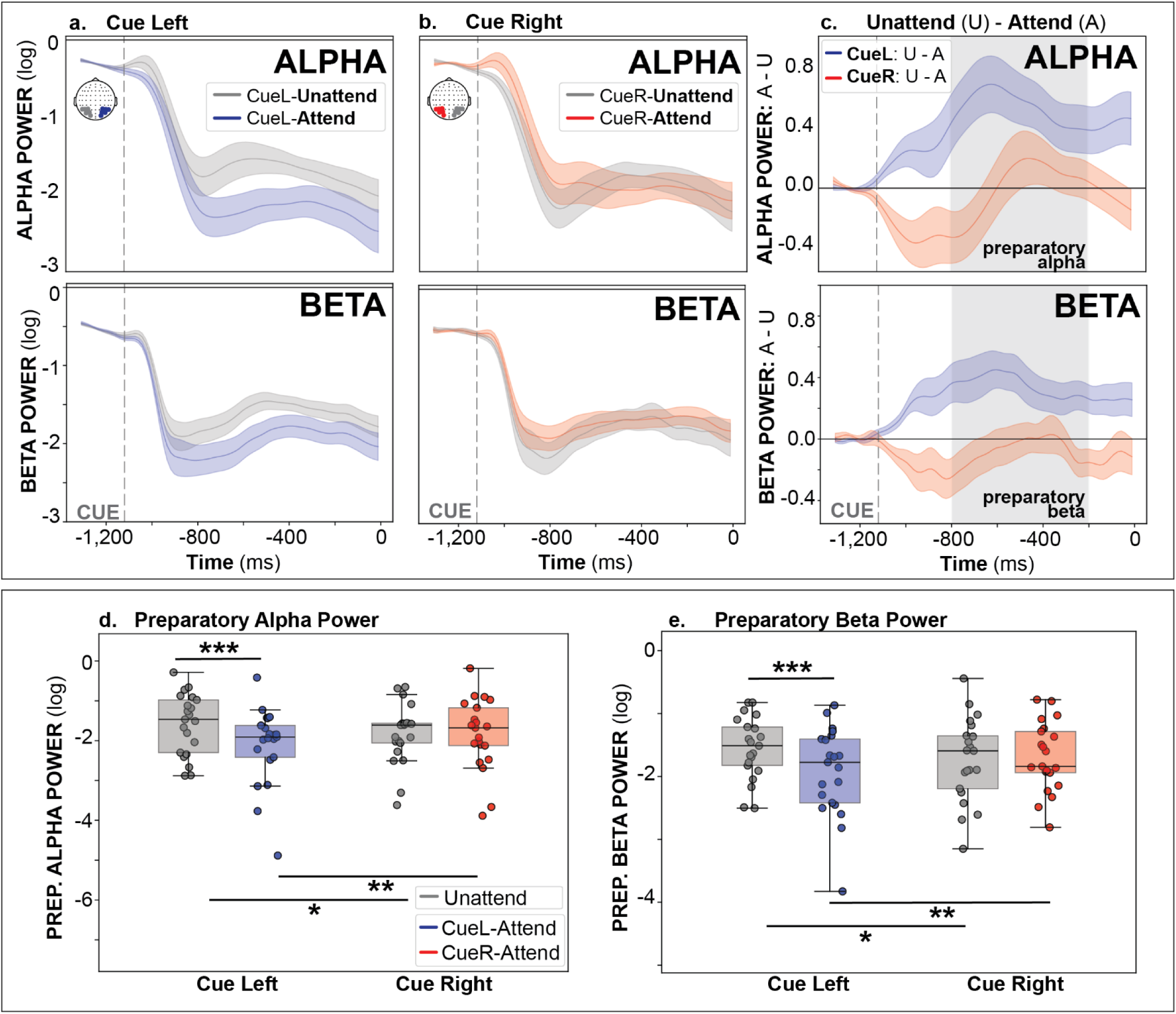
Attentional modulation of alpha and beta-power differs by cue side. **a-b)** Time courses of log-transformed alpha-band (top; 7-13-Hz; posterior electrodes) and beta-band (bottom; 13-20-Hz; posterior electrodes) power aligned to cue onset for Cue Left **(a)** and Cue Right **(b)** trials. Attended (blue: cue-left; red: cue-right) and unattended (gray) traces are overlaid, with shaded regions representing ±SEM. Attended responses corresponds to the ROI contralateral to the cued hemifield, and unattended corresponds to the ROI ipsilateral to the cued hemifield.The time course represents the post-cue window prior to S2, in which the dashed vertical line marks cue onset. **c)** Attention effects are shown as the difference wave between attended and unattended ROI (A-U) for both Cue Left (blue) and Cue Right (red). The preparatory region, indexing preparation for stimuli, is highlighted by the shaded gray time window (−800 to −200 ms). **d-e)** Boxplots beneath each ERP panel summarize mean alpha **(d)** and beta **(e)** power, separated by group and cue-side. Asterisks indicate significant effects of attention (attended vs. unattended ROI) or cue side (left vs. right) (*p < 0.05, **p < 0.01, ***p < 0.001).

Parallel effects were observed in beta-band power (**Fig. 5e**), with significant cue-direction differences in both regions of interest, such that power over the unattended ROI was greater for left versus right attention (unattended: left – right β = 0.182, p = 0.012) while power over the attended ROI showed the opposite pattern (attended: left – right β = −0.228, p = 0.002). Again, power over the unattended ROI also exceeded power over the attended ROI for leftward-directed attention (left: unattended – attended. β = 0.367, p < 0.001).

A significant main effect of Block was also observed in the alpha-band (F = 6.722, p = 0.0011), reflecting lower alpha power during inter-mixed blocks compared to pure social (IM – S: β = −0.295, p = 0.002) and pure non-social (IM – NS: β = −0.247, p = 0.013) blocks. Similar effects were observed in the beta band (F = 3.167, p = 0.044), with a marginal, non-significant reduction in beta power during inter-mixed blocks relative to pure social blocks (IM – S: β = −0.123, p = 0.121) and pure non-social blocks (IM – NS: β = −0.146, p = 0.053).

No significant effects of Age were observed for alpha power (p>0.05), but older age was associated with lower beta power (β = −0.063, p = 0.049).

### Resting Alpha Power

In an analysis of alpha power at rest (**Supplementary Fig. 2**, **Supplementary Table 1**), LMMs revealed a main effect of Condition (*F* = 52.395, *p* < 0.001), with higher mean power in the eyes-closed (vs. eyes-open) condition (open − closed: B = −0.387, *p* < 0.001). A similar effect was observed for the aperiodic exponent, with a significant main effect of Condition (F = 38.652, p < 0.001), reflecting higher exponent values in the eyes-closed relative to eyes-open condition (open − closed: β = −0.151, p < 0.001). This steeper spectral slope is consistent with a shift toward greater inhibitory dominance (i.e., a lower E/I index) during eyes-closed rest.

There was also a main effect of RS-Block (*F* = 17.65, *p* < 0.001), reflecting an overall increase in mean power in the resting state power collected before-task versus after-task (before − after: B = 0.215, *p* < 0.001). A comparable main effect of RS-Block was also observed for the aperiodic exponent (F = 24.787, p < 0.001), with higher exponent values before relative to after the task (before − after: β = 0.121, p < 0.001), consistent with a shift toward a more inhibition-dominant (lower E/I) state before task. A similar main effect of RS-Block was observed for alpha peak frequency (F = 8.439, p = 0.004), with higher frequencies before relative to after the task (before − after: β = 0.267, p = 0.004), indicating a modest pre-task shift toward faster intrinsic alpha dynamics.

These effects were qualified by a significant Condition × RS-Block interaction (*F* = 5.453, *p* = 0.021), indicating that task-related changes in mean power differed as a function of eyes-open vs. eyes-closed conditions. Follow-up contrasts showed that mean power was lower in the open relative to closed condition both before (before: open – closed. B = −0.507, *p* < 0.001) and after (after: open – closed. B = −0.268, *p* = 0.001). Examination of task-related changes within each condition revealed mean alpha power was significantly higher pre-task versus post-task in the eyes-closed condition (eyes-closed: before – after. B = 0.334, *p* < 0.001), whereas no significant pre–post task change was observed in the eyes-open condition (eyes-open: before – after. B = 0.095, *p* = 0.19).

Age was significantly associated with spectral parameters at rest, such that alpha peak frequency increased with age (β = 0.103, p = 0.018), while the aperiodic exponent decreased with age (β = −0.03, p = 0.002). There was no significant effect of age on mean power.

Critically, we found no significant differences in alpha power, frequency or aperiodic exponent over left versus right parieto-occipital regions at rest.

### Theta phase–RT relationships and cross-frequency power coupling

To further disentangle the mechanisms and drivers underlying spatial attention in this task—and particularly the observed cue-side asymmetries—we next examined anticipatory theta phase dynamics during the cue-S2 interval, which revealed a significant relationship between fronto-central theta phase (ipsilateral to cue-direction) and RT. As shown in **Fig. 6a-b**, topographical mapping of phase–RT modulation revealed significant effects over FC1 during cue-left trials at 3-Hz (FFT-amplitude = 1.677; p = 0.017), and over FC2 during cue-right trials at 6-Hz (FFT-amplitude = 1.708; p = 0.014) and 7-Hz (FFT-amplitude = 1.535; p = 0.035). In both cases, ipsilateral-cue theta phase accounted for systematic fluctuations in behavioral performance, with specific phase angles (“good” phase) corresponding to faster RTs.

**Figure 6.**
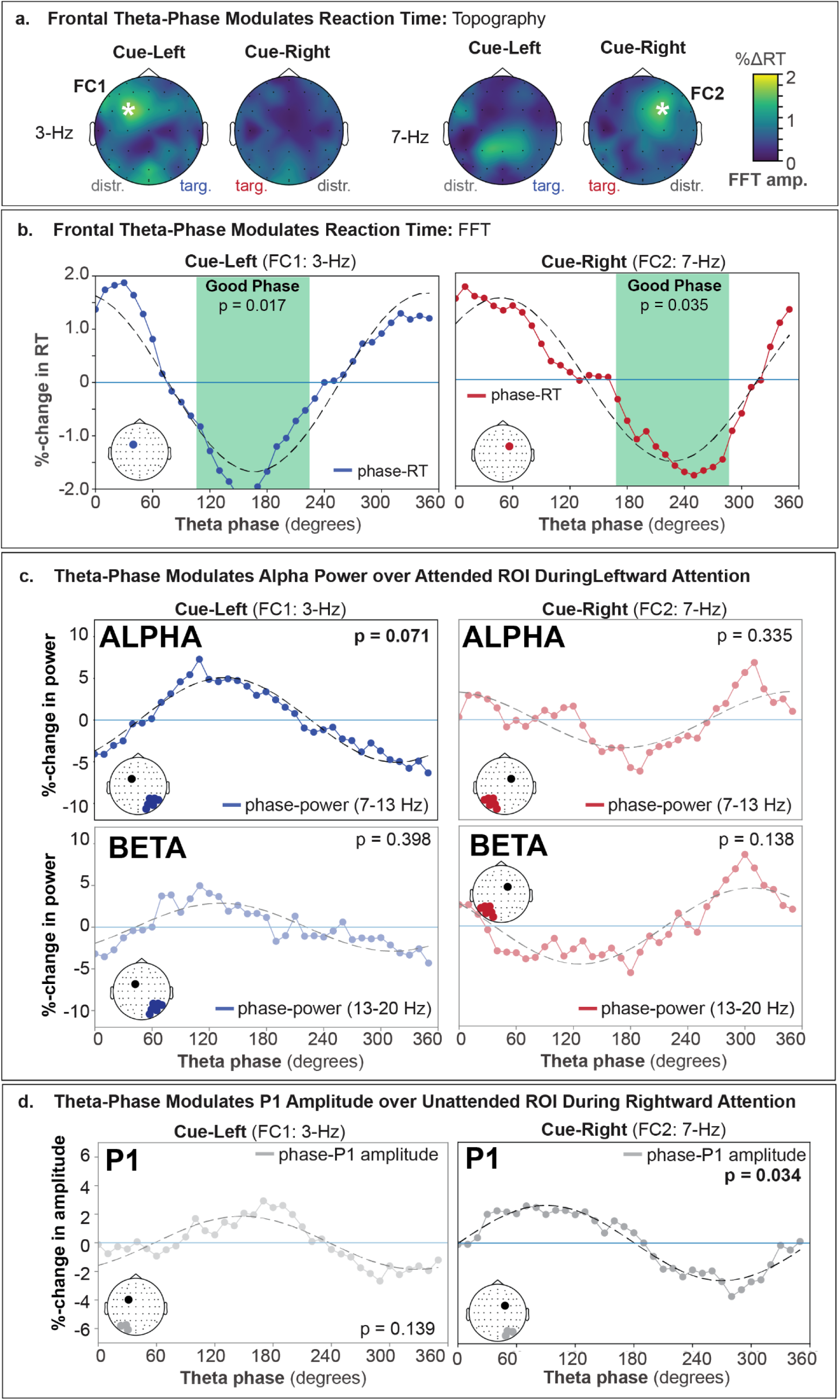
Ipsilateral-cue frontal theta phase–dependent modulation of RT, parieto-occipital neuro-oscillatory power and P1 amplitude. **a)** Scalp topographies showing the spatial distribution of the FFT amplitude of the anticipatory theta phase–reaction-time (RT) relationship, separately for cue-left and cue-right trials. For cue-left trials, a significant effect is observed at 3-Hz over left fronto-central cortex (FC1), whereas for cue-right trials, the effect is observed at 7-Hz over right fronto-central cortex (FC2). Color scale reflects FFT amplitude of percent change in RT (%ΔRT); asterisks denote electrodes showing statistically significant phase–RT modulation (permutation test, p < 0.05). A significant phase–behavior relationship was also detected at 6-Hz over FC2 for cue-right attention, although we present only the 7-Hz results for clarity and brevity, as the 6-Hz effect demonstrated equivalent phase–behavior and phase-power coupling at this site. **b)** Theta phase–RT functions extracted from the fronto-central electrodes and frequencies identified in panel (a): cue-left trials at FC1 (3-Hz: left) and cue-right trials at FC2 (7-Hz: right). Blue (cue-left) and red (cue-right) traces show mean percent change in RT across theta phase bins (0–360°), with negative values indicating faster responses. Black dashed lines represent the best-fitting 1-cycle FFT. Shaded green regions indicate the phase window associated with maximal behavioral facilitation (“good” phase), defined as the contiguous phase range around the RT minimum. Reported p-values reflect the significance of the 1-cycle FFT amplitude derived from permutation testing. **c)** Theta phase–power functions with phase extracted from fronto-central electrodes at the frequencies identified in panel (a): cue-left trials at FC1 (3-Hz; left) and cue-right trials at FC2 (7-Hz, right). Power was measured from parieto-occipital channels contralateral to the to-be-attended space (right parieto-occipital for cue-left; left parieto-occipital for cue-right) at the same time points used for instantaneous phase extraction for alpha- (7-13-Hz; top) and beta- (13-20 Hz; bottom) bands. Blue (cue-left) and red (cue-right) traces show mean percent change in power across theta phase bins (0–360°), with negative values indicating reduced power. Black dashed lines denote the best-fitting one-cycle FFT. Reported p-values reflect the significance of the one-cycle phase–power FFT amplitude derived from permutation testing, indicating a phase-dependent modulation of power. **d)** Theta phase–P1 functions with phase extracted from fronto-central electrodes at the frequencies identified in panel (a): cue-left trials at FC1 (3-Hz; left) and cue-right trials at FC2 (7-Hz, right). Mean P1 amplitude (110-150 ms) over the unattended ROI was measured from parieto-occipital ERP channels contralateral to the to-be-ignored (unattended) location (left parieto-occipital for cue-left; right parieto-occipital for cue-right). Blue (cue-left) and red (cue-right) traces show mean percent change in P1 amplitude across theta phase bins (0–360°), with negative values indicating a reduction in P1 amplitude. Black dashed lines denote the best-fitting one-cycle FFT. Reported p-values reflect the significance of the one-cycle phase–power FFT amplitude derived from permutation testing, indicating a phase-dependent modulation of P1 amplitude.

To assess whether these behaviorally optimal theta phases were associated with concurrent modulation of sensory processing, we examined if identified theta-phase (cue-left: 3-Hz at FC1; cue-right: 6-7-Hz at FC2) correlated with parieto-occipital alpha and beta power (**Fig. 6c**). A marginal association was observed selectively during cue-left trials in the alpha band (7–13-Hz; FFT amplitude = 5.093; p = 0.071) over the attended (right hemisphere) parieto-occipital ROI. Given the possibility that this effect reflected a narrower spectral contribution within the alpha band, we next examined individual frequencies (7–13-Hz; **Supplementary Fig. 3**). This analysis revealed that the association was driven specifically by 9-Hz (FFT amplitude = 5.418; p = 0.048) and 10-Hz (FFT amplitude = 5.301; p = 0.047) power over the attended (right hemisphere) parieto-occipital region during cue-left trials. No other alpha frequencies showed significant phase–power coupling.

No significant phase–power relationships were observed over the unattended ROI for alpha, for either hemisphere for beta frequencies, or during cue-right trials (ps>0.05).

However, because attentional modulation of alpha and beta modulation over both attended and unattended ROIs remained evident in primary power analyses, we considered whether contralateral (to cue) frontal theta phase might contribute to this effect, even in the absence of a behavioral relationship. We therefore examined whether contralateral-cue frontal theta (cue-left: 3-Hz at FC2; cue-right: 6–7-Hz at FC1) was associated with modulation of parieto-occipital alpha and beta power. Although alpha power (over attended and unattended ROI) was not significantly modulated by contralateral-cue theta phase, we observed a significant association between contralateral (right-hemisphere) frontal theta phase and beta power over the unattended (left hemisphere) ROI (13–20 Hz; FFT amplitude = 6.3431; p = 0.0406) during cue-left trials (**Supplementary Fig. 4**). We did not identify a significant relationship between contralateral-cue theta phase and alpha or beta modulation of either hemisphere during cue-right trials.

Because we observed robust sensory gain effects—indexed by attentional modulation of P1 amplitude—for both leftward and rightward attention, we next tested whether ipsilateral frontal theta phase also modulated P1 amplitude in each condition (**Fig. 6d**). This relationship emerged selectively during rightward attention. Specifically, 7-Hz (FFT amplitude = 2.621, p = 0.034)—but not 6-Hz (FFT amplitude = 1.667, p = 0.263)—frontal theta modulated P1 amplitude over the unattended ROI. In contrast, no significant theta–P1 coupling was observed during leftward attention (FFT amplitude = 1.861, p = 0.139). Theta-phase did not significantly modulate P1 amplitude corresponding to attended ROI in leftward (3-Hz: FFT amplitude = 0.065, p = 0.694) or rightward attention (6-Hz: FFT amplitude = 1.501, p = 0.275; 7-Hz: FFT amplitude = 2.081, p = 0.076).

Together, these findings support a model of spatial attention in which ipsilateral fronto-central theta phase organizes behavioral performance in a cue-direction–specific manner, with distinct frequencies governing leftward (3-Hz) and rightward (6–7-Hz) attention. Critically, theta-phase coupling to sensory gain was asymmetric. During leftward attention, theta phase was selectively associated with alpha modulation over the attended ROI, whereas during rightward attention, theta phase correlated with modulation of early sensory processing, as indexed by P1 amplitude. In contrast, contralateral theta phase, which exhibited no behavioral coupling, was linked to beta modulation over the unattended ROI only during leftward attention.

To determine whether pre-stimulus alpha power or early sensory gain (P1 amplitude) more strongly predicted behavioral performance—and whether these effects were driven by attended versus unattended regions—we conducted linear mixed-effects modeling of trial-level RT (RT) with subject-level random intercepts (**Figure 7**).

**Figure 7.**
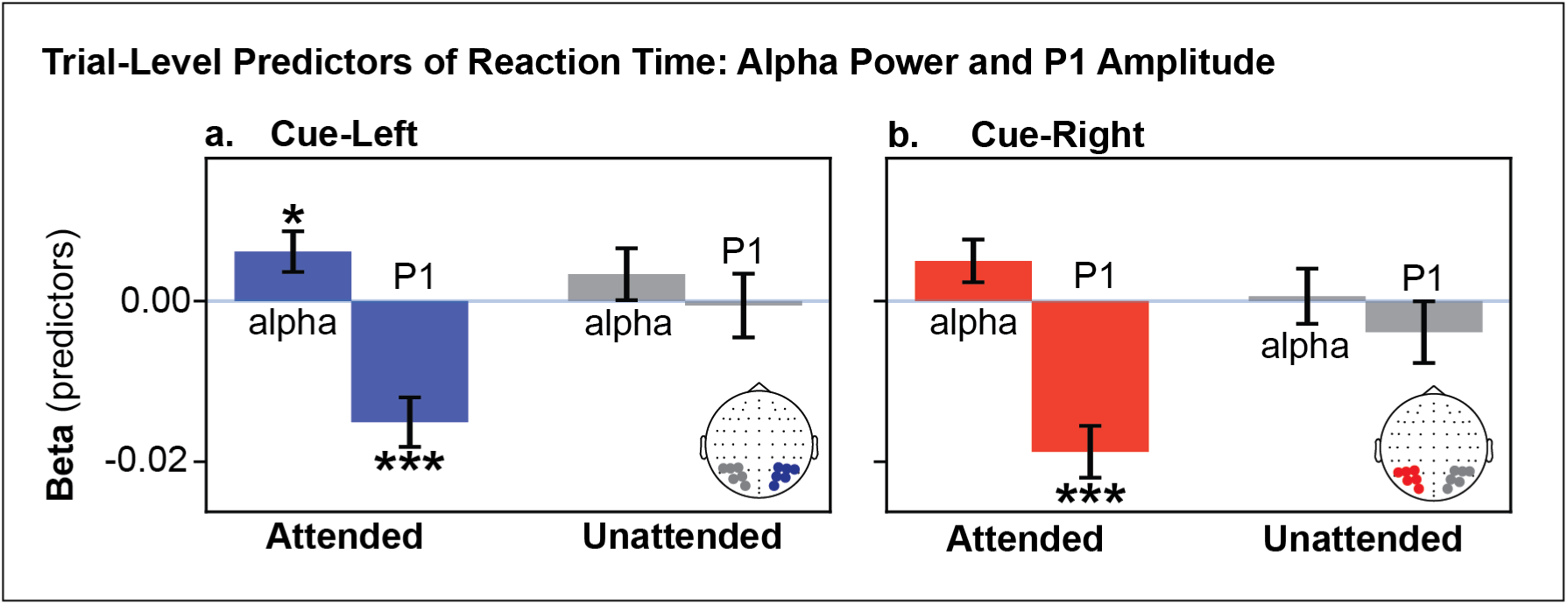
Trial-level oscillatory and sensory predictors of RT. Standardized beta coefficients (± SE) from LMM estimating trial-level RT (RT) as a function of pre-stimulus parieto-occipital alpha power (−800 to −200 ms prior to S2 onset) and P1 amplitude, shown separately for cue-left **(a)** and cue-right **(b)** trials and for attended (cue-left: right hemisphere; cue-right: left hemisphere) and unattended (cue-left: left hemisphere; cue-right: right hemisphere) ROI. Positive beta values indicate longer RTs with increasing predictor magnitude, whereas negative beta values indicate faster RTs. Models included subject-level random intercepts to account for within-subject trial clustering. Significance thresholds: * p < .05, ** p < .01, *** p < .001.

Critically, significant effects were confined to the ROI corresponding to the to-be-attended space. For cue-left trials, lower alpha power over the attended ROI was associated with faster RTs (β = 0.0062, SE = 0.0025, *p* = 0.015), indicating that enhanced alpha desynchronization over the attended ROI predicted quicker target detection. In contrast, larger P1 amplitudes over the attended ROI predicted faster RTs (β = −0.0151, SE = 0.0031, *p* < 0.001), consistent with enhanced early sensory gain facilitating behavioral performance.

For cue-right trials, P1 amplitude remained a robust predictor of RT (β = −0.0187, SE = 0.0032, *p* < 0.001), whereas the alpha effect was weaker and did not reach significance (β = 0.0050, SE = 0.0027, *p* = .060). Thus, while P1 sensory gain modulated RT across cue directions, alpha modulation of behavior was evident primarily during cue-left trials.

Importantly, no significant effects emerged for alpha or P1 amplitude over the unattended ROI, indicating that behavioral variability was driven specifically by modulation of the a to-be-attended space rather than suppression of the to-be-ignored space.

To confirm that the behavioral modulation observed during leftward attention reflected coordinated sensory–oscillatory dynamics, we next examined whether P1 amplitude covaried with pre-stimulus alpha power at the trial level (**Supplementary Figure 5**). Consistent with the mixed-effects results, we observed a significant association between pre-stimulus alpha power and subsequent P1 amplitude over the attended ROI during cue-left trials (r = 0.027, p = 0.004), such that lower alpha states were associated with larger P1 responses. In contrast, this relationship was not observed during cue-right trials (r = 0.008, p = 0.428).

Given the observed asymmetry in alpha-related behavioral modulation during leftward relative to rightward attention, we next examined whether this hemispheric difference varied across development. Preparatory alpha modulation asymmetry increased significantly with age (B = 0.173, p = 0.021; **Fig. 8**), indicating that older participants exhibited stronger hemispheric differentiation in anticipatory alpha power. Specifically, the left–right difference in unattended–attended alpha modulation became more positive across development, reflecting increasingly greater relative alpha modulation during leftward compared to rightward attention. These findings suggest that hemispheric asymmetry in anticipatory sensory gating strengthens with age.

**Figure 8.**
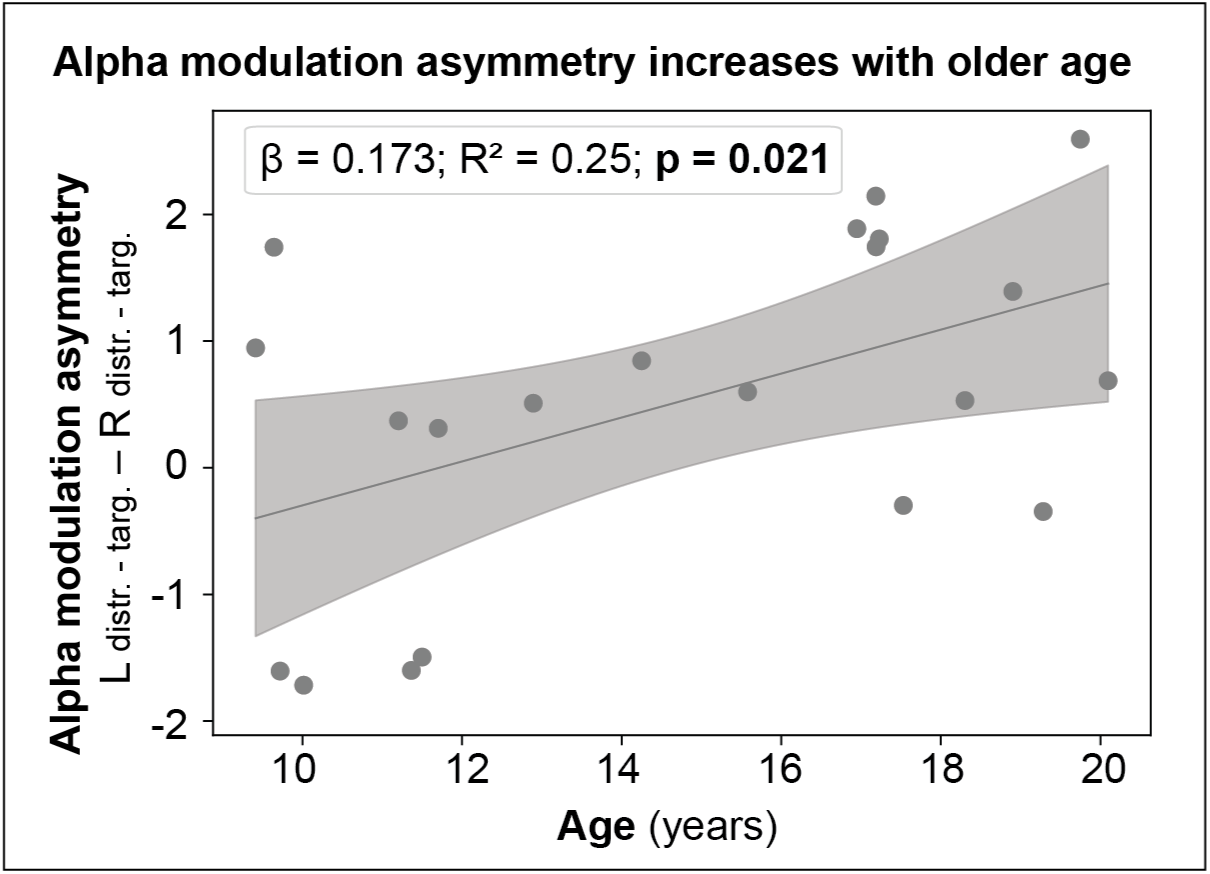
Developmental increases in asymmetric alpha modulation during the anticipatory window. Relationship between age and preparatory alpha modulation asymmetry. Alpha modulation asymmetry was quantified as the left–right difference in unattended–attended alpha power (log-transformed) computed from parieto-occipital electrodes during the −800 to −200 ms pre-stimulus interval, indexing hemispheric differences in anticipatory sensory gating. Positive values reflect stronger relative alpha modulation for leftward-directed attention compared with rightward-directed attention. Each point represents an individual participant; the solid line denotes the best-fitting linear regression, with shaded regions indicating the 95% confidence interval.

## Discussion

Together, the findings presented here provide strong support for rhythmic theories of attention, indicating that attentional control is temporally structured by ongoing oscillatory phase rather than continuously deployed. Intrinsic frontal theta phase predicted trial-by-trial fluctuations in behavioral performance, demonstrating that endogenous oscillatory dynamics can organize perceptual sensitivity. Extending prior work that has primarily relied on externally imposed phase resets, we show that intrinsic theta oscillatory rhythms asymmetrically structure attentional control in the developing human brain, providing mechanistic insight into the well-described phenomenon of pseudoneglect. Finally, this work establishes a developmental framework in which frontal theta coordinates posterior excitation to promote selective spatial attention, leveraging a multimodal approach that integrates high-density electrophysiology (EEG), pupillometry, and cognitive measures within the same participants.

### Top-down frontal theta orchestrates behavior

The present findings support a model in which frontal theta phase provides a temporally structured control signal that shapes moment-to-moment fluctuations in spatial attention. Across cue directions, fronto-central theta phase predicted systematic fluctuations in RT, consistent with the idea that attentional control is not continuously deployed but instead cycles through phases of relative facilitation versus relative suppression.

To our knowledge, this is the first demonstration that intrinsic frontal theta phase reliably predicts behavioral performance in a human EEG paradigm using a fixed interstimulus interval that permits precise alignment of phase to target onset. Prior studies examining phase–behavior relationships have typically relied on manipulations that perturb or reset oscillatory phase, such as jittering the interstimulus interval, rhythmic sensory stimulation (e.g., flashes of light), or transcranial magnetic stimulation (Landau and Fries 2012, Fiebelkorn, Saalmann et al. 2013, Landau, Schreyer et al. 2015, Fiebelkorn, Pinsk et al. 2018, Helfrich, Fiebelkorn et al. 2018, Gordon, Belardinelli et al. 2022, Plochl, Fiebelkorn et al. 2022, Cavanah and Fiebelkorn 2025, Lo, Ke et al. 2025, Trajkovic, Veniero et al. 2025, Han, Brincat et al. 2026). In contrast, our paradigm used a fixed cue–S2 interval without phase-resetting manipulations, demonstrating that theta-phase–dependent fluctuations in behavior can emerge from ongoing endogenous neural dynamics.

Moreover, prior work on the role of theta modulation of attention has involved only adults. This is the first evidence that such theta-phase–dependent modulation of behavior can be robustly observed across development. By demonstrating that anticipatory frontal theta phase systematically organizes RT in a developing population, the present findings extend rhythmic models of attention into a developmental framework and establish intrinsic theta dynamics as a measurable and behaviorally relevant control mechanism across maturation.

An additional and unexpected feature of the present results is that the behaviorally relevant theta signal was consistently observed over the hemisphere ipsilateral to the cue—that is, over the frontal site corresponding to the to-be-ignored (unattended) space—during both leftward and rightward attention. Notably, this spatial pattern echoes prior neuroimaging findings showing that direction of attention modulates frontal recruitment asymmetrically, with leftward attention engaging right occipital and left frontal regions more strongly, and rightward attention engaging right frontal cortex (Vandenberghe, Duncan et al. 1997, Vandenberghe, Duncan et al. 2000). At present, the functional significance of this ipsilateral-cue localization remains unclear. One possibility is that frontal theta reflects a control process engaged in suppressing or stabilizing the unattended hemifield, rather than directly enhancing the attended representation. Alternatively, ipsilateral frontal cortex may coordinate interhemispheric dynamics, indirectly biasing posterior sensory regions through competitive or supervisory control mechanisms. Because this pattern was consistent across cue directions and replicated in both behavioral and cross-frequency analyses, it is unlikely to be spurious; however, its mechanistic basis remains unresolved. Future work incorporating source modeling and connectivity or causal perturbation approaches will be needed to determine whether ipsilateral frontal theta indexes distractor suppression, interhemispheric competition, or a broader supervisory control signal.

### Behavior is selectively modulated by sensory gain in the attended ROI

A central contribution of the present work is the dissociation between enhancement of the attended space versus suppression of the unattended space as determinants of behavioral performance. Although anticipatory alpha and beta dynamics showed robust attention-related lateralization, consistent with preparatory cortical gating, trial-level predictors of RT were confined to the region corresponding to the to-be-attended space. Larger P1 amplitudes predicted faster responses for both cue directions, establishing early sensory gain as the most proximal neural mechanism to behavior in this task. In contrast, P1 amplitude and alpha power over the unattended ROI did not explain trial-to-trial RT variability, despite clear evidence that these signals were modulated by attention at the group level. This dissociation indicates that, under the demands of the present paradigm, behavioral variability is governed predominantly by how effectively the system modulates the attended representation rather than how strongly it suppresses the unattended representation.

Importantly, the direction of alpha modulation further supports this interpretation. During the anticipatory window, alpha power over the attended region did not increase above the fixation baseline; rather, attention was characterized by relative alpha desynchronization. This pattern is more consistent with an excitability-enhancing mechanism—facilitating target identification through sustained desynchronization (Mulholland 1965, Pfurtscheller 1992, Bastiaansen and Brunia 2001, Sauseng, Klimesch et al. 2005)—than with a purely inhibitory gating model (Kelly, Lalor et al. 2006), a heavily debated framework in the field. These results do not argue against an active suppressive role for alpha, which is strongly supported in sustained, high-competition paradigms (e.g. (Kelly, Lalor et al. 2006, Banerjee, Snyder et al. 2011, Murphy, Foxe et al. 2014)); rather, they indicate that in the present speeded-detection context, trial-to-trial RT variability is most strongly governed by sensory gain over the attended region.

The relationship between anticipatory alpha dynamics and early sensory responses remains a central point of debate in the attention literature. One influential account proposes that the P1 is simply a manifestation of ongoing alpha phase—essentially a traveling wave of alpha oscillatory activity (Gruber, Klimesch et al. 2005, Klimesch, Hanslmayr et al. 2007, Fellinger, Klimesch et al. 2011, Klimesch, Fellinger et al. 2011, Trajkovic, Di Gregorio et al. 2024). In contrast, other work argues that alpha activity and the P1 reflect functionally dissociable mechanisms (Slagter, Prinssen et al. 2016). In this context, we posit that a reduction in alpha reflects a release from inhibition that renders the attended cortex more responsive, thereby enabling larger P1 amplitudes and faster detection (Tarasi, di Pellegrino et al. 2022). The fact that lower pre-stimulus alpha predicted both larger P1 amplitudes (during cue-left trials) and faster RTs reinforces the interpretation that alpha desynchronization operates as a permissive mechanism for early sensory gain.

### Specialized attentional mechanisms: Cue-side asymmetries in theta control of posterior alpha/beta drive a leftward attentional bias

The asymmetric oscillatory architecture observed here provides a potential mechanistic account for the well-known phenomenon of pseudoneglect, that is, the subtle but reliable leftward bias in spatial attention observed in healthy individuals. Here, we observed stronger alpha-mediated suppression during leftward attention, consistent with models proposing dominant right-hemisphere control of spatial attention, which may require coordinated inhibitory mechanisms to bias sensory processing toward the left visual field. In contrast, rightward attention may rely on more distributed bilateral control, reducing the need for strong alpha-mediated suppression. A representative schematic illustrating these proposed mechanisms is shown in **Fig. 9**.

**Figure 9.**
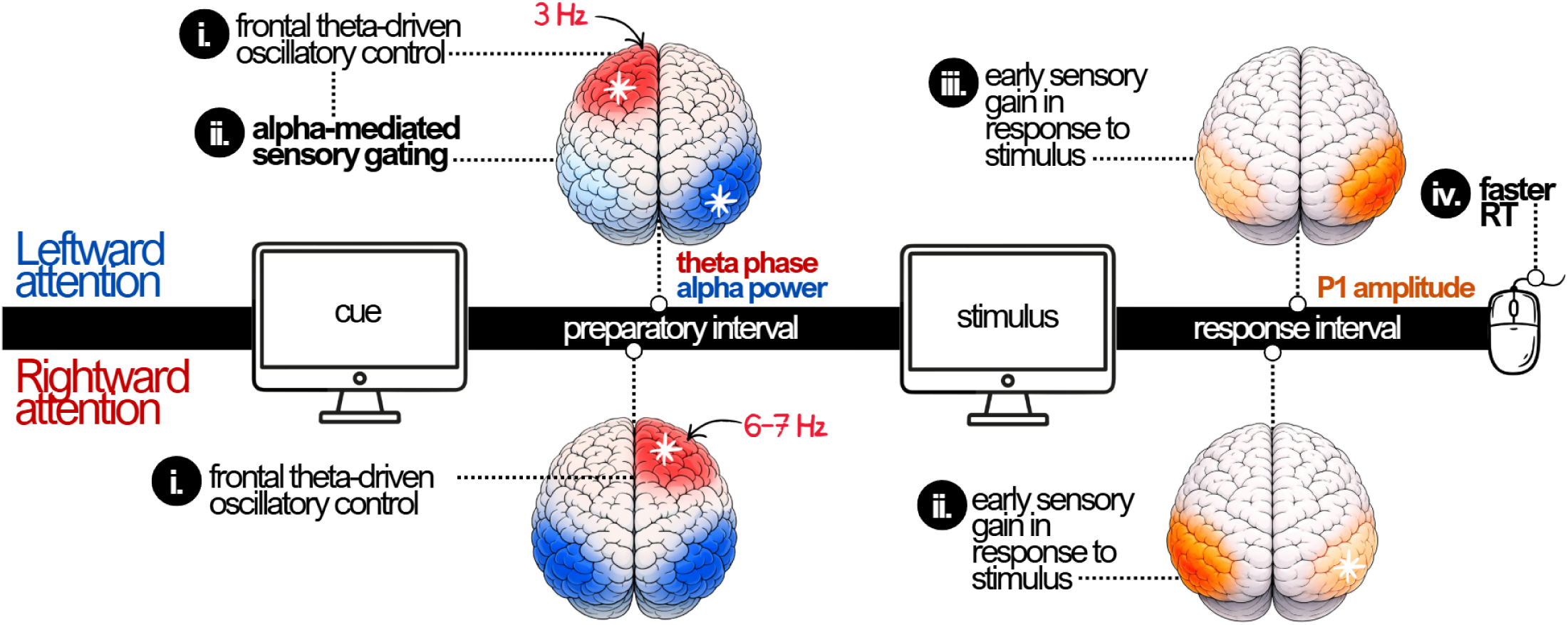
Asymmetric oscillatory processes supporting rightward and leftward spatial attention. Schematic illustrating a model of the neural mechanisms underlying leftward (top) and rightward (bottom) spatial attention. Following presentation of a spatial cue, activity during the anticipatory interval is characterized by frontal theta–driven control signals that are ipsilateral to cued space, with theta phase linked to behavioral performance. During leftward attention, frontal theta (∼3-Hz) coordinates alpha-mediated sensory gating across parieto-occipital regions, in which theta phase is linked to changes in alpha power over the right hemisphere. This preparatory alpha modulation facilitates early sensory gain upon stimulus onset, reflected in enhanced P1 amplitude in attended posterior visual regions and faster behavioral responses relative to rightward attention. During rightward attention, the relationship between frontal theta-phase and behavior occurs at a higher frequency (∼6-7-Hz) and is not accompanied by posterior alpha modulation. Instead, theta activity modulates P1 amplitude, leading to early sensory modulation of the P1 component.

Behaviorally, participants responded faster when attention was cued to the left, demonstrating a leftward advantage that was not accompanied by differences in hit rate, false alarms, inverse efficiency score (IES), or d′. This pattern is consistent with longstanding evidence that the human attentional system is not bilaterally symmetric, with relative facilitation for leftward attention often attributed to right-hemisphere dominance in spatial attention (Bradshaw, Nathan et al. 1987, McCourt and Jewell 1999, Orr and Nicholls 2005, Charles, Sahraie et al. 2007, Sosa, Teder-Salejarvi et al. 2010). The present findings extend this literature by providing a temporally resolved account of how this asymmetry may be implemented at the level of oscillatory control and sensory gain. During leftward (cue-left) attention, we observed a significant phase-behavior relationship between frontal 3-Hz theta over the hemisphere ipsilateral to the cue (FC1). This rhythm predicted RT and exhibited phase-dependent coordination with posterior oscillatory dynamics. Specifically, theta phase modulated alpha power (9–10-Hz within the alpha band) over the attended ROI and beta power over the unattended ROI. At the trial level, RT was predicted by both alpha desynchronization and P1 amplitude in the attended ROI, and these two measures were significantly correlated. Lower pre-stimulus alpha was associated with larger P1 amplitudes, and both lower alpha and larger P1 amplitudes predicted faster responses. These relationships are consistent with a cascade in which frontal theta organizes posterior excitability states via alpha modulation, shaping early sensory gain (P1), which in turn relates to behavioral speed. Frontal theta–posterior alpha coupling has been reported previously in adults and non-human primates (Klimesch, Sauseng et al. 2007, Klimesch, Sauseng et al. 2007, Romei, Gross et al. 2010, Hanslmayr, Gross et al. 2011, Wilson and Foxe 2020, Han, Brincat et al. 2026), but only one prior study (Sauseng, Klimesch et al. 2005) directly compared coupling strength between attended and unattended regions, demonstrating stronger modulation of alpha power corresponding to the attended (vs. unattended) region. However, they did not directly test how alpha modulation varies between leftward and rightward attention and did not incorporate eye tracking to verify sustained covert attention.

In contrast, rightward (cue-right) attention was characterized by a higher theta frequency (6–7-Hz) over ipsilateral frontal cortex (FC2). Although theta phase robustly predicted RT, it did not significantly correlate with posterior alpha or beta power over either hemisphere. At the behavioral level, RT during rightward trials was predicted by P1 amplitude alone, with no significant contribution from pre-stimulus alpha. Moreover, frontal theta phase modulated P1 amplitude only during rightward attention. These findings suggest a different control architecture in which frontal theta relates to early sensory gain, without evidence for coordinated anticipatory alpha-mediated gating in posterior cortex. This interpretation is broadly consistent with recent reports showing that pre-stimulus theta phase can modulate the amplitude of stimulus-evoked ERP components, although those studies typically collapse across cue direction (Cavanah and Fiebelkorn 2026, Redding, Ding et al. 2026).

Taken together, the data indicate that the leftward behavioral advantage is associated with an oscillatory control profile in which slower frontal theta relates to posterior alpha modulation and sensory gain. Rightward attention, in contrast, appears to involve a faster frontal rhythm that relates to early sensory responses without recruitment of oscillatory modulation. Importantly, resting-state alpha did not differ between hemispheres, indicating that these asymmetries do not simply arise from baseline hemispheric alpha power differences, but instead emerge during task-driven spatial control.

Pupillometry and sensory ERP latency findings further contextualize these effects. Increased stimulus-evoked pupil responses during rightward attention support the thesis that rightward orienting requires greater attentional engagement. Pupil dilation has been shown to track cognitive effort required to deploy covert spatial attention (Brocher, Harbecke et al. 2018), and recent work combining pupillometry and fMRI demonstrates greater dilation during attend-right relative to attend-left cues, suggesting that rightward orienting may require additional top-down control to overcome the typical left visual field bias (Meyyappan, Rajan et al. 2023). Given that pupil dynamics have been linked to locus-coeruleus–noradrenergic (LC-NE) modulation of attentional state (Joshi, Li et al. 2016, Wang, Baird et al. 2018, Joshi and Gold 2020, Joshi 2021), these findings suggest that the asymmetric oscillatory dynamics observed here may operate within a broader neuromodulatory framework. Recent evidence suggests that LC-NE activity enhances perceptual sensitivity (Ghosh and Maunsell 2024), raising the possibility that larger pupil responses during rightward orienting may reflect greater neuromodulatory support for sensory processing, particularly when oscillatory preparatory engagement is weaker. In addition, P1 latency was delayed for stimuli in the right visual field relative to the left visual field, which may suggest a greater attentional load during rightward orienting (Fu, Fedota et al. 2010). Together, the EEG and pupil findings here suggest that distinct control architectures support attention to the left and right visual fields. However, the precise relationship between arousal, oscillatory control, and sensory timing remains to be determined.

This interpretation is consistent with prior electrophysiological findings. Thut and colleagues demonstrated an asymmetric bias in posterior alpha, with greater modulation during leftward compared to rightward attention, which corresponded to enhanced leftward behavioral performance (Thut, Nietzel et al. 2006). Extending this work, the present results suggest that hemispheric asymmetries in spatial attention reflect differences in temporal control architecture—specifically in theta-mediated top-down coordination—rather than simple differences in oscillatory power. These conclusions also align with neuroimaging evidence of asymmetric frontoparietal recruitment during spatial attention. Siman-Tov and colleagues (2007) demonstrated stronger and more bilateral frontoparietal engagement during leftward attention, linked to enhanced intraparietal sulcus connectivity, whereas rightward attention was characterized by more unilateral recruitment (Siman-Tov, Mendelsohn et al. 2007).

An important remaining question, however, concerns the functional significance of the frequency difference between leftward and rightward attention. Specifically, leftward attention was associated with slower frontal theta (3-Hz), whereas rightward attention engaged a faster rhythm (6–7-Hz). One possibility is that slower theta during leftward attention reflects more sustained control states, facilitating coordinated modulation of posterior oscillatory activity. In contrast, faster theta during rightward attention may reflect a more rapid control regime, potentially consistent with prior work linking higher alpha and theta frequencies to faster rhythmic sampling (Busch, Dubois et al. 2009, Dugue, Marque et al. 2011, Kienitz, Schmiedt et al. 2018, Yang, Yuan et al. 2025). Alternatively, these differences may reflect hemispheric specialization within frontal control networks, although future work will be necessary to adjudicate among these possibilities.

### Developmental strengthening of leftward–rightward asymmetry in spatial attentional control

A consistent developmental signature emerged across measures. As expected, older participants responded faster, made fewer false alarms, showed higher discrimination performance (d′), and demonstrated improved efficiency, indicating broad maturation of attentional performance. At the neural level, age was associated with earlier P1 latencies and reduced P1 amplitudes, suggesting developmental shifts toward faster, potentially more efficient early sensory processing. At rest, alpha peak frequency increased while the aperiodic exponent decreased with age, consistent with developmental changes in background spectral organization.

Crucially, the asymmetry in anticipatory alpha modulation strengthened with age: older participants exhibited greater left–right differentiation in unattended–attended alpha modulation over posterior regions, reflecting increasing lateralization of preparatory sensory gating across development. One interpretation is that with maturation, the neural architecture supporting lateralized control—potentially involving frontoparietal networks that bias posterior cortex—becomes more differentiated, enabling more distinct preparatory states depending on the cued direction of attention. Alternatively, this may reflect a maturational shift toward more strongly lateralized attentional states. Pseudoneglect has been reported in healthy children (Michel, Bidot et al. 2011), but is significantly understudied and findings are highly conflicting (see (Kaul, Papadatou-Pastou et al. 2023) for review). Notably, the relationship between pseudoneglect and age remains inconsistent: some work reports increased pseudoneglect with advancing age (Chiffi, Diana et al. 2021), whereas other studies show a reduction in hemispheric asymmetry in older populations (Casagrande, Agostini et al. 2021). However, these studies examine age-related changes extending into elderly populations, leaving asymmetries in attention relatively uncharacterized in a developmental context.

In this context, the fact that this developmental increase in asymmetry appears in task-evoked preparatory dynamics but not in resting hemispheric differences supports the view that development enhanced the deployment and coordination of posterior oscillations such that they become more lateralized. In combination with the leftward-specific coupling between preparatory alpha and sensory gain, these developmental findings imply that the mechanisms linking control to early sensory processing become more directionally specialized across development, potentially contributing to increasingly stable and efficient attentional selection into adulthood.

### Limitations

Although the age-related effects were consistent across behavioral, sensory, and spectral measures, the sample size was modest and spanned a broad developmental range. Accordingly, inferences about maturational trajectories should be considered preliminary and would benefit from replication in larger, age-stratified cohorts.

A second consideration concerns motor lateralization, as the task required right-handed motor responses in a sample of predominantly right-handed individuals. As described by the Simon effect (Valle-Inclan 1996, Proctor, Zhong et al. 2023), motor preparation and effector dominance could, in principle, interact with hemispheric control dynamics and influence oscillatory or sensory measures. Importantly, however, P1 and anticipatory alpha analyses (**Fig. 4-5**) were conducted exclusively on non-target trials, thereby eliminating motor execution and response preparation as confounds in the neural measures. Moreover, the behavioral leftward advantage we observed would not be predicted by a simple Simon compatibility account: in right-handed participants, the Simon effect would instead predict a relative response advantage for trials in the compatible (right-side) visual hemi-field. The fact that our behavioral asymmetry aligns with pseudoneglect rather than response-side compatibility argues against a purely motoric explanation. Nevertheless, future studies incorporating counterbalanced response hands and mixed-handed samples will be important to fully dissociate attentional asymmetries from potential motor-related lateralization effects.

## Author Contributions

M.D and S.M conceived the study. M.D was responsible for participant recruitment and data collection and curation. M.D pre-processed and analyzed the data under the supervision of T.V, C.B and S.M. M.D wrote the first draft of the manuscript. S.M, J.J.F, T.V and C.B provided edits on the manuscript. All authors approved the final manuscript.

## List of Abbreviations

CPT-3: Conners’ Continuous Performance Test, 3rd Edition
d′: Detection sensitivity (d-prime)
EEG: Electroencephalography
ERP: Event-related potential
FA: False alarm
FFT: Fast Fourier Transform
FS-IQ: Full-scale intelligence quotient
Hz: Hertz
ICA: Independent component analysis
IES: Inverse efficiency score
IM: Inter-mixed (block condition)
LMM: Linear mixed-effects model
LSL: Lab Streaming Layer
NS: Non-social
PO: Parieto-occipital
P1: First positive visual ERP component (∼100 ms)
ROI: Region of interest
RT: Reaction time
RS: Resting state
S: Social
SEM: Standard error of the mean
SOA: Stimulus-onset asynchrony
SRS-2: Social Responsiveness Scale, 2nd Edition
S1: Stimulus 1 (cue)
S2: Stimulus 2 (stimuli/target display)
TD: Typically developing
UDTR: Up-Down Transformed Rule
V-IQ: Verbal intelligence quotient

## Acknowledgements & Funding

Support for recruitment and phenotyping of participants was provided by the Human Clinical Phenotyping Core of the NICHD funded Rose F. Kennedy Intellectual and Developmental Disabilities Research Center (P50 HD105352). Work at the collaborating site in Rochester is supported through the B. Thomas Golisano Intellectual and Developmental Disabilities Research Institute (UR-IDDRC), which is supported by a center grant from the Eunice Kennedy Shriver National Institute of Child Health and Human Development (P50 HD103536 – to JJF). Partial support was also provided by the Albert Einstein College of Medicine Medical Scientist Training Program (T32-GM149364).

## References

Awh, E., L. Anllo-Vento and S. A. Hillyard (2000). "The role of spatial selective attention in working memory for locations: evidence from event-related potentials." J Cogn Neurosci 12(5): 840–847.

Banerjee, S., A. C. Snyder, S. Molholm and J. J. Foxe (2011). "Oscillatory alpha-band mechanisms and the deployment of spatial attention to anticipated auditory and visual target locations: supramodal or sensory-specific control mechanisms?" J Neurosci 31(27): 9923–9932.

Bartolomeo, P. and S. Chokron (2002). "Orienting of attention in left unilateral neglect." Neurosci Biobehav Rev 26(2): 217–234.

Basar, E., M. Schurmann, C. Basar-Eroglu and S. Karakas (1997). "Alpha oscillations in brain functioning: an integrative theory." Int J Psychophysiol 26(1-3): 5–29.

Bastiaansen, M. C. and C. H. Brunia (2001). "Anticipatory attention: an event-related desynchronization approach." Int J Psychophysiol 43(1): 91–107.

Bates D, M. M., Bolker B, Walker S (2015). "Fitting Linear Mixed-Effects Models Using lme4." Journal of Statistical Software 67(1): 1–48.

Berger, H. (1929). "Über das Elektrenkephalogramm des Menschen." Archiv f. Psychiatrie 87: 527–570.

Bigdely-Shamlo, N., T. Mullen, C. Kothe, K. M. Su and K. A. Robbins (2015). "The PREP pipeline: standardized preprocessing for large-scale EEG analysis." Front Neuroinform 9: 16.

Bonnefond, M. and O. Jensen (2025). "The role of alpha oscillations in resisting distraction." Trends Cogn Sci 29(4): 368–379.

Bradshaw, J. L., G. Nathan, N. C. Nettleton, L. Wilson and J. Pierson (1987). "Why is there a left side underestimation in rod bisection?" Neuropsychologia 25(4): 735–738.

Brandt, M. E. and B. H. Jansen (1991). "The relationship between prestimulus-alpha amplitude and visual evoked potential amplitude." Int J Neurosci 61(3-4): 261–268.

Brandt, M. E., B. H. Jansen and J. P. Carbonari (1991). "Pre-stimulus spectral EEG patterns and the visual evoked response." Electroencephalogr Clin Neurophysiol 80(1): 16–20.

Brocher, A., R. Harbecke, T. Graf, D. Memmert and S. Huttermann (2018). "Using task effort and pupil size to track covert shifts of visual attention independently of a pupillary light reflex." Behav Res Methods 50(6): 2551–2567.

Busch, N. A., J. Dubois and R. VanRullen (2009). "The phase of ongoing EEG oscillations predicts visual perception." J Neurosci 29(24): 7869–7876.

Canolty, R. T., E. Edwards, S. S. Dalal, M. Soltani, S. S. Nagarajan, H. E. Kirsch, M. S. Berger, N. M. Barbaro and R. T. Knight (2006). "High gamma power is phase-locked to theta oscillations in human neocortex." Science 313(5793): 1626–1628.

Casagrande, M., F. Agostini, F. Favieri, G. Forte, J. Giovannoli, A. Guarino, A. Marotta, F. Doricchi and D. Martella (2021). "Age-Related Changes in Hemispherical Specialization for Attentional Networks." Brain Sci 11(9).

Cavanah, P. J. and I. C. Fiebelkorn (2025). "A shared theta-rhythmic process for selective sampling of environmental information and internally stored information." bioRxiv.

Cavanah, P. J. and I. C. Fiebelkorn (2026). "A shared theta-rhythmic process for selective sampling of environmental information and internally stored information." J Neurosci.

Charles, J., A. Sahraie and P. McGeorge (2007). "Hemispatial asymmetries in judgment of stimulus size." Percept Psychophys 69(5): 687–698.

Chiffi, K., L. Diana, M. Hartmann, D. Cazzoli, C. L. Bassetti, R. M. Muri and A. K. Eberhard-Moscicka (2021). "Spatial asymmetries ("pseudoneglect") in free visual exploration-modulation of age and relationship to line bisection." Exp Brain Res 239(9): 2693–2700.

Chokron, S. (2003). "Right parietal lesions, unilateral spatial neglect, and the egocentric frame of reference." Neuroimage 20 **Suppl 1**: S75–81.

Colby, C. L. (1991). "The neuroanatomy and neurophysiology of attention." J Child Neurol 6 **Suppl**: S90–118.

Conners, C. K. (2008). "Conners third edition (Conners 3)." Los Angeles, CA: Western Psychological Services.

Constantino, J. N., S. A. Davis, R. D. Todd, M. K. Schindler, M. M. Gross, S. L. Brophy, L. M. Metzger, C. S. Shoushtari, R. Splinter and W. Reich (2003). "Validation of a brief quantitative measure of autistic traits: comparison of the social responsiveness scale with the autism diagnostic interview-revised." J Autism Dev Disord 33(4): 427–433.

Corbetta, M. and G. L. Shulman (2011). "Spatial neglect and attention networks." Annu Rev Neurosci 34: 569–599.

Desimone, R. and J. Duncan (1995). "Neural mechanisms of selective visual attention." Annu Rev Neurosci 18: 193–222.

Donoghue, T., M. Haller, E. J. Peterson, P. Varma, P. Sebastian, R. Gao, T. Noto, A. H. Lara, J. D. Wallis, R. T. Knight, A. Shestyuk and B. Voytek (2020). "Parameterizing neural power spectra into periodic and aperiodic components." Nat Neurosci 23(12): 1655–1665.

Dugue, L., P. Marque and R. VanRullen (2011). "The phase of ongoing oscillations mediates the causal relation between brain excitation and visual perception." J Neurosci 31(33): 11889–11893.

Dugue, L., P. Marque and R. VanRullen (2015). "Theta oscillations modulate attentional search performance periodically." J Cogn Neurosci 27(5): 945–958.

E Basar, M. S. (1996). Alpha Rhythms in the Brain: Functional Correlates. Physiology. 11: 57–102.

Fellinger, R., W. Klimesch, W. Gruber, R. Freunberger and M. Doppelmayr (2011). "Pre-stimulus alpha phase-alignment predicts P1-amplitude." Brain Res Bull 85(6): 417–423.

Fiebelkorn, I. C. and S. Kastner (2019). "A Rhythmic Theory of Attention." Trends Cogn Sci 23(2): 87–101.

Fiebelkorn, I. C., M. A. Pinsk and S. Kastner (2018). "A Dynamic Interplay within the Frontoparietal Network Underlies Rhythmic Spatial Attention." Neuron 99(4): 842–853 e848.

Fiebelkorn, I. C., Y. B. Saalmann and S. Kastner (2013). "Rhythmic sampling within and between objects despite sustained attention at a cued location." Curr Biol 23(24): 2553–2558.

Fiebelkorn, I. C., A. C. Snyder, M. R. Mercier, J. S. Butler, S. Molholm and J. J. Foxe (2013). "Cortical cross-frequency coupling predicts perceptual outcomes." Neuroimage 69: 126–137.

Foxe, J. J., M. E. McCourt and D. C. Javitt (2003). "Right hemisphere control of visuospatial attention: line-bisection judgments evaluated with high-density electrical mapping and source analysis." Neuroimage 19(3): 710–726.

Foxe, J. J., G. V. Simpson and S. P. Ahlfors (1998). "Parieto-occipital approximately 10 Hz activity reflects anticipatory state of visual attention mechanisms." Neuroreport 9(17): 3929–3933.

Foxe, J. J. and A. C. Snyder (2011). "The Role of Alpha-Band Brain Oscillations as a Sensory Suppression Mechanism during Selective Attention." Front Psychol 2: 154.

Frey, H. P., A. M. Schmid, J. W. Murphy, S. Molholm, E. C. Lalor and J. J. Foxe (2014). "Modulation of early cortical processing during divided attention to non-contiguous locations." Eur J Neurosci 39(9): 1499–1507.

Frischen, A., A. P. Bayliss and S. P. Tipper (2007). "Gaze cueing of attention: visual attention, social cognition, and individual differences." Psychol Bull 133(4): 694–724.

Frund, I., N. A. Busch, J. Schadow, U. Korner and C. S. Herrmann (2007). "From perception to action: phase-locked gamma oscillations correlate with reaction times in a speeded response task." BMC Neurosci 8: 27.

Fu, K. M., J. J. Foxe, M. M. Murray, B. A. Higgins, D. C. Javitt and C. E. Schroeder (2001). "Attention-dependent suppression of distracter visual input can be cross-modally cued as indexed by anticipatory parieto-occipital alpha-band oscillations." Brain Res Cogn Brain Res 12(1): 145–152.

Fu, S., J. R. Fedota, P. M. Greenwood and R. Parasuraman (2010). "Dissociation of visual C1 and P1 components as a function of attentional load: an event-related potential study." Biol Psychol 85(1): 171–178.

Ghosh, S. and J. H. R. Maunsell (2024). "Locus coeruleus norepinephrine contributes to visual-spatial attention by selectively enhancing perceptual sensitivity." Neuron 112(13): 2231–2240 e2235.

Gomez-Ramirez, M., S. P. Kelly, S. Molholm, P. Sehatpour, T. H. Schwartz and J. J. Foxe (2011). "Oscillatory sensory selection mechanisms during intersensory attention to rhythmic auditory and visual inputs: a human electrocorticographic investigation." J Neurosci 31(50): 18556–18567.

Gordon, P. C., P. Belardinelli, M. Stenroos, U. Ziemann and C. Zrenner (2022). "Prefrontal theta phase-dependent rTMS-induced plasticity of cortical and behavioral responses in human cortex." Brain Stimul 15(2): 391–402.

Gramfort, A., M. Luessi, E. Larson, D. A. Engemann, D. Strohmeier, C. Brodbeck, R. Goj, M. Jas, T. Brooks, L. Parkkonen and M. Hamalainen (2013). "MEG and EEG data analysis with MNE-Python." Front Neurosci 7: 267.

Gray, C. M., P. Konig, A. K. Engel and W. Singer (1989). "Oscillatory responses in cat visual cortex exhibit inter-columnar synchronization which reflects global stimulus properties." Nature 338(6213): 334–337.

Gray, C. M. and W. Singer (1989). "Stimulus-specific neuronal oscillations in orientation columns of cat visual cortex." Proc Natl Acad Sci U S A 86(5): 1698–1702.

Gruber, T., M. M. Muller, A. Keil and T. Elbert (1999). "Selective visual-spatial attention alters induced gamma band responses in the human EEG." Clin Neurophysiol 110(12): 2074–2085.

Gruber, W. R., W. Klimesch, P. Sauseng and M. Doppelmayr (2005). "Alpha phase synchronization predicts P1 and N1 latency and amplitude size." Cereb Cortex 15(4): 371–377.

Guillery, S. M. S. R. W. (2001). Exploring the Thalamus. Amsterdam, Elsevier.

Han, H. B., S. L. Brincat, T. J. Buschman and E. K. Miller (2026). "Working memory readout varies with frontal theta rhythms." Neuron 114(1): 159–166 e152.

Hanslmayr, S., J. Gross, W. Klimesch and K. L. Shapiro (2011). "The role of alpha oscillations in temporal attention." Brain Res Rev 67(1-2): 331–343.

Heilman, K. M., E. Valenstein and R. T. Watson (2000). "Neglect and related disorders." Semin Neurol 20(4): 463–470.

Heilman, K. M. and T. Van Den Abell (1980). "Right hemisphere dominance for attention: the mechanism underlying hemispheric asymmetries of inattention (neglect)." Neurology 30(3): 327–330.

Heinze, H. J., G. R. Mangun, W. Burchert, H. Hinrichs, M. Scholz, T. F. Munte, A. Gos, M. Scherg, S. Johannes, H. Hundeshagen and et al. (1994). "Combined spatial and temporal imaging of brain activity during visual selective attention in humans." Nature 372(6506): 543–546.

Helfrich, R. F., I. C. Fiebelkorn, S. M. Szczepanski, J. J. Lin, J. Parvizi, R. T. Knight and S. Kastner (2018). "Neural Mechanisms of Sustained Attention Are Rhythmic." Neuron 99(4): 854–865 e855.

Hu, F. and Y. Dan (2022). "An inferior-superior colliculus circuit controls auditory cue-directed visual spatial attention." Neuron 110(1): 109–119 e103.

J.N., C. (2012). "Social Responsiveness Scale (2nd ed.)." WPS Publish.

James, W. (1890). The Principles of Psychology, Henry Holt and Co.

Jas, M., D. A. Engemann, Y. Bekhti, F. Raimondo and A. Gramfort (2017). "Autoreject: Automated artifact rejection for MEG and EEG data." Neuroimage 159: 417–429.

Joshi, S. (2021). "Pupillometry: Arousal State or State of Mind?" Curr Biol 31(1): R32–R34.

Joshi, S. and J. I. Gold (2020). "Pupil Size as a Window on Neural Substrates of Cognition." Trends Cogn Sci 24(6): 466–480.

Joshi, S., Y. Li, R. M. Kalwani and J. I. Gold (2016). "Relationships between Pupil Diameter and Neuronal Activity in the Locus Coeruleus, Colliculi, and Cingulate Cortex." Neuron 89(1): 221–234.

Kaul, D., M. Papadatou-Pastou and G. Learmonth (2023). "A meta-analysis of the line bisection task in children." Laterality 28(1): 48–71.

Keller, A. S., I. Davidesco and K. D. Tanner (2020). "Attention Matters: How Orchestrating Attention May Relate to Classroom Learning." CBE Life Sci Educ 19(3): fe5.

Kelly, S. P., M. Gomez-Ramirez and J. J. Foxe (2009). "The strength of anticipatory spatial biasing predicts target discrimination at attended locations: a high-density EEG study." Eur J Neurosci 30(11): 2224–2234.

Kelly, S. P., E. C. Lalor, R. B. Reilly and J. J. Foxe (2006). "Increases in alpha oscillatory power reflect an active retinotopic mechanism for distracter suppression during sustained visuospatial attention." J Neurophysiol 95(6): 3844–3851.

Kienitz, R., J. T. Schmiedt, K. A. Shapcott, K. Kouroupaki, R. C. Saunders and M. C. Schmid (2018). "Theta Rhythmic Neuronal Activity and Reaction Times Arising from Cortical Receptive Field Interactions during Distributed Attention." Curr Biol 28(15): 2377–2387 e2375.

Klimesch, W., R. Fellinger and R. Freunberger (2011). "Alpha oscillations and early stages of visual encoding." Front Psychol 2: 118.

Klimesch, W., S. Hanslmayr, P. Sauseng, W. R. Gruber and M. Doppelmayr (2007). "P1 and traveling alpha waves: evidence for evoked oscillations." J Neurophysiol 97(2): 1311–1318.

Klimesch, W., P. Sauseng and S. Hanslmayr (2007). "EEG alpha oscillations: the inhibition-timing hypothesis." Brain Res Rev 53(1): 63–88.

Klimesch, W., P. Sauseng, S. Hanslmayr, W. Gruber and R. Freunberger (2007). "Event-related phase reorganization may explain evoked neural dynamics." Neurosci Biobehav Rev 31(7): 1003–1016.

Kothe, C., S. Y. Shirazi, T. Stenner, D. Medine, C. Boulay, M. I. Grivich, T. Mullen, A. Delorme and S. Makeig (2024). "The Lab Streaming Layer for Synchronized Multimodal Recording." bioRxiv.

Krajbich, I., C. Armel and A. Rangel (2010). "Visual fixations and the computation and comparison of value in simple choice." Nat Neurosci 13(10): 1292–1298.

Krajbich, I. and A. Rangel (2011). "Multialternative drift-diffusion model predicts the relationship between visual fixations and choice in value-based decisions." Proc Natl Acad Sci U S A 108(33): 13852–13857.

Krauzlis, R. J., L. P. Lovejoy and A. Zenon (2013). "Superior colliculus and visual spatial attention." Annu Rev Neurosci 36: 165–182.

Landau, A. N. and P. Fries (2012). "Attention samples stimuli rhythmically." Curr Biol 22(11): 1000–1004.

Landau, A. N., H. M. Schreyer, S. van Pelt and P. Fries (2015). "Distributed Attention Is Implemented through Theta-Rhythmic Gamma Modulation." Curr Biol 25(17): 2332–2337.

Lenth, M. R., Singmann, H., Love, J., Buerkner, P. & Herve, M (2019). "Package ‘emmeans’."

Lo, Y. H., S. C. Ke, T. T. Chen, H. H. Tsao and P. Tseng (2025). "Theta burst entrainment of human EEG using flickering light stimulation." J Neurophysiol 134(5): 1466–1475.

Luck, S. J., H. J. Heinze, G. R. Mangun and S. A. Hillyard (1990). "Visual event-related potentials index focused attention within bilateral stimulus arrays. II. Functional dissociation of P1 and N1 components." Electroencephalogr Clin Neurophysiol 75(6): 528–542.

Maris, E. and R. Oostenveld (2007). "Nonparametric statistical testing of EEG- and MEG-data." J Neurosci Methods 164(1): 177–190.

Martin A. Fischler, R. C. B. (1981). "Random sample consensus: a paradigm for model fitting with applications to image analysis and automated cartography." Communications of the ACM 24(6): 381–395.

Mathot, S. (2018). "Pupillometry: Psychology, Physiology, and Function." J Cogn 1(1): 16.

Mathot, S. and A. Vilotijevic (2023). "Methods in cognitive pupillometry: Design, preprocessing, and statistical analysis." Behav Res Methods 55(6): 3055–3077.

McCourt, M. E. and G. Jewell (1999). "Visuospatial attention in line bisection: stimulus modulation of pseudoneglect." Neuropsychologia 37(7): 843–855.

Meyyappan, S., A. Rajan, G. R. Mangun and M. Ding (2023). "Top-down control of the left visual field bias in cued visual spatial attention." Cereb Cortex 33(9): 5097–5107.

Michel, C., S. Bidot, F. Bonnetblanc and P. Quercia (2011). "Left minineglect or inverse pseudoneglect in children with dyslexia?" Neuroreport 22(2): 93–96.

Miller, E. K. and T. J. Buschman (2013). "Cortical circuits for the control of attention." Curr Opin Neurobiol 23(2): 216–222.

Mulholland, T. B. (1965). "Occurrence of the electroencephalographic alpha rhythm with eyes open." Nature 206(985): 746.

Murphy, J. W., J. J. Foxe and S. Molholm (2016). "Neuro-oscillatory mechanisms of intersensory selective attention and task switching in school-aged children, adolescents and young adults." Dev Sci 19(3): 469–487.

Murphy, J. W., J. J. Foxe, J. B. Peters and S. Molholm (2014). "Susceptibility to distraction in autism spectrum disorder: probing the integrity of oscillatory alpha-band suppression mechanisms." Autism Res 7(4): 442–458.

Nummenmaa, L. and A. J. Calder (2009). "Neural mechanisms of social attention." Trends Cogn Sci 13(3): 135–143.

Orr, C. A. and M. E. Nicholls (2005). "The nature and contribution of space- and object-based attentional biases to free-viewing perceptual asymmetries." Exp Brain Res 162(3): 384–393.

Perkovic, S., M. Schoemann, C. J. Lagerkvist and J. L. Orquin (2023). "Covert attention leads to fast and accurate decision-making." J Exp Psychol Appl 29(1): 78–94.

Pernet, C. R., M. Latinus, T. E. Nichols and G. A. Rousselet (2015). "Cluster-based computational methods for mass univariate analyses of event-related brain potentials/fields: A simulation study." J Neurosci Methods 250: 85–93.

Perrin, F., J. Pernier, O. Bertrand and J. F. Echallier (1989). "Spherical splines for scalp potential and current density mapping." Electroencephalogr Clin Neurophysiol 72(2): 184–187.

Petersen, S. E., M. Corbetta, F. M. Miezin and G. L. Shulman (1994). "PET studies of parietal involvement in spatial attention: comparison of different task types." Can J Exp Psychol 48(2): 319–338.

Petersen, S. E., D. L. Robinson and J. D. Morris (1987). "Contributions of the pulvinar to visual spatial attention." Neuropsychologia 25(1A): 97–105.

Pfurtscheller, G. (1992). "Event-related synchronization (ERS): an electrophysiological correlate of cortical areas at rest." Electroencephalogr Clin Neurophysiol 83(1): 62–69.

Plochl, M., I. Fiebelkorn, S. Kastner and J. Obleser (2022). "Attentional sampling of visual and auditory objects is captured by theta-modulated neural activity." Eur J Neurosci 55(11-12): 3067–3082.

Proctor, R. W., Q. Zhong and J. Chen (2023). "The Simon Effect Asymmetry for Left- and Right-Dominant Persons." J Cogn 6(1): 17.

Redding, Z. V., Y. Ding and I. C. Fiebelkorn (2026). "Frequency-specific attentional mechanisms phasically modulate the influence of distractors on task performance." PLoS Biol 24(2): e3003664.

Rich, E. L. and J. D. Wallis (2017). "Spatiotemporal dynamics of information encoding revealed in orbitofrontal high-gamma." Nat Commun 8(1): 1139.

Romei, V., J. Gross and G. Thut (2010). "On the role of prestimulus alpha rhythms over occipito-parietal areas in visual input regulation: correlation or causation?" J Neurosci 30(25): 8692–8697.

Sauseng, P., W. Klimesch, W. Stadler, M. Schabus, M. Doppelmayr, S. Hanslmayr, W. R. Gruber and N. Birbaumer (2005). "A shift of visual spatial attention is selectively associated with human EEG alpha activity." Eur J Neurosci 22(11): 2917–2926.

Sherman, S. M. (2001). "Tonic and burst firing: dual modes of thalamocortical relay." Trends Neurosci 24(2): 122–126.

Siman-Tov, T., A. Mendelsohn, T. Schonberg, G. Avidan, I. Podlipsky, L. Pessoa, N. Gadoth, L. G. Ungerleider and T. Hendler (2007). "Bihemispheric leftward bias in a visuospatial attention-related network." J Neurosci 27(42): 11271–11278.

Slagter, H. A., S. Prinssen, L. C. Reteig and A. Mazaheri (2016). "Facilitation and inhibition in attention: Functional dissociation of pre-stimulus alpha activity, P1, and N1 components." Neuroimage 125: 25–35.

Snyder, A. C. and J. J. Foxe (2010). "Anticipatory attentional suppression of visual features indexed by oscillatory alpha-band power increases: a high-density electrical mapping study." J Neurosci 30(11): 4024–4032.

Sosa, Y., W. A. Teder-Salejarvi and M. E. McCourt (2010). "Biases of spatial attention in vision and audition." Brain Cogn 73(3): 229–235.

Spagna, A., T. H. Kim, T. Wu and J. Fan (2020). "Right hemisphere superiority for executive control of attention." Cortex 122: 263–276.

Stone, S. P., P. W. Halligan and R. J. Greenwood (1993). "The incidence of neglect phenomena and related disorders in patients with an acute right or left hemisphere stroke." Age Ageing 22(1): 46–52.

Szczepanski, S. M., N. E. Crone, R. A. Kuperman, K. I. Auguste, J. Parvizi and R. T. Knight (2014). "Dynamic changes in phase-amplitude coupling facilitate spatial attention control in fronto-parietal cortex." PLoS Biol 12(8): e1001936.

Szczepanski, S. M., C. S. Konen and S. Kastner (2010). "Mechanisms of spatial attention control in frontal and parietal cortex." J Neurosci 30(1): 148–160.

Tarasi, L., G. di Pellegrino and V. Romei (2022). "Are you an empiricist or a believer? Neural signatures of predictive strategies in humans." Prog Neurobiol 219: 102367.

Thakral, P. P. and S. D. Slotnick (2009). "The role of parietal cortex during sustained visual spatial attention." Brain Res 1302: 157–166.

Thut, G., A. Nietzel, S. A. Brandt and A. Pascual-Leone (2006). "Alpha-band electroencephalographic activity over occipital cortex indexes visuospatial attention bias and predicts visual target detection." J Neurosci 26(37): 9494–9502.

Tort, A. B., R. W. Komorowski, J. R. Manns, N. J. Kopell and H. Eichenbaum (2009). "Theta-gamma coupling increases during the learning of item-context associations." Proc Natl Acad Sci U S A 106(49): 20942–20947.

Townsend, J. and E. Courchesne (1994). "Parietal damage and narrow "spotlight" spatial attention." J Cogn Neurosci 6(3): 220–232.

Trajkovic, J., F. Di Gregorio, G. Thut and V. Romei (2024). "Transcranial magnetic stimulation effects support an oscillatory model of ERP genesis." Curr Biol 34(5): 1048–1058 e1044.

Trajkovic, J., D. Veniero, S. Hanslmayr, S. Palva, G. Cruz, V. Romei and G. Thut (2025). "Top-down and bottom-up interactions rely on nested brain oscillations to shape rhythmic visual attention sampling." PLoS Biol 23(4): e3002688.

Tsuchimoto, S., S. Shibusawa, S. Iwama, M. Hayashi, K. Okuyama, N. Mizuguchi, K. Kato and J. Ushiba (2021). "Use of common average reference and large-Laplacian spatial-filters enhances EEG signal-to-noise ratios in intrinsic sensorimotor activity." J Neurosci Methods 353: 109089.

Valle-Inclan, F. (1996). "The locus of interference in the Simon effect: an ERP study." Biol Psychol 43(2): 147–162.

Vandenberghe, R., J. Duncan, K. M. Arnell, S. J. Bishop, N. J. Herrod, A. M. Owen, P. S. Minhas, P. Dupont, J. D. Pickard and G. A. Orban (2000). "Maintaining and shifting attention within left or right hemifield." Cereb Cortex 10(7): 706–713.

Vandenberghe, R., J. Duncan, P. Dupont, R. Ward, J. B. Poline, G. Bormans, J. Michiels, L. Mortelmans and G. A. Orban (1997). "Attention to one or two features in left or right visual field: a positron emission tomography study." J Neurosci 17(10): 3739–3750.

Wang, C. A., T. Baird, J. Huang, J. D. Coutinho, D. C. Brien and D. P. Munoz (2018). "Arousal Effects on Pupil Size, Heart Rate, and Skin Conductance in an Emotional Face Task." Front Neurol 9: 1029.

Watson, B. O., M. Ding and G. Buzsaki (2018). "Temporal coupling of field potentials and action potentials in the neocortex." Eur J Neurosci 48(7): 2482–2497.

Weschler, D. (2011). "Wechsler Abbreviated Scale of Intelligence (2nd ed.)." Pearson.

Weschler, D. (2014). "Wechsler Intelligence Scale for Children (5th ed.)." Pearson.

Wetherill, G. B. and H. Levitt (1965). "Sequential Estimation of Points on a Psychometric Function." Br J Math Stat Psychol 18: 1–10.

Wijers, A. A., J. J. Lange, G. Mulder and L. J. Mulder (1997). "An ERP study of visual spatial attention and letter target detection for isoluminant and nonisoluminant stimuli." Psychophysiology 34(5): 553–565.

Wilson, T. J. and J. J. Foxe (2020). "Cross-frequency coupling of alpha oscillatory power to the entrainment rhythm of a spatially attended input stream." Cogn Neurosci 11(1-2): 71–91.

Woldorff, M. G., P. T. Fox, M. Matzke, J. L. Lancaster, S. Veeraswamy, F. Zamarripa, M. Seabolt, T. Glass, J. H. Gao, C. C. Martin and P. Jerabek (1997). "Retinotopic organization of early visual spatial attention effects as revealed by PET and ERPs." Hum Brain Mapp 5(4): 280–286.

Worden, M. S., J. J. Foxe, N. Wang and G. V. Simpson (2000). "Anticipatory biasing of visuospatial attention indexed by retinotopically specific alpha-band electroencephalography increases over occipital cortex." J Neurosci 20(6): RC63.

Yang, F., P. Yuan, L. Shen, K. Zhou, S. He and Y. Jiang (2025). "Visual awareness sharpens and accelerates attentional sampling through enhancing inhibitory neural modulation in the attention network." Nat Commun 16(1): 10058.

Zhou, H., R. J. Schafer and R. Desimone (2016). "Pulvinar-Cortex Interactions in Vision and Attention." Neuron 89(1): 209–220.

