## Supplementary Materials for "Frontal theta phase modulates asymmetric posterior neural mechanisms of spatial attention"

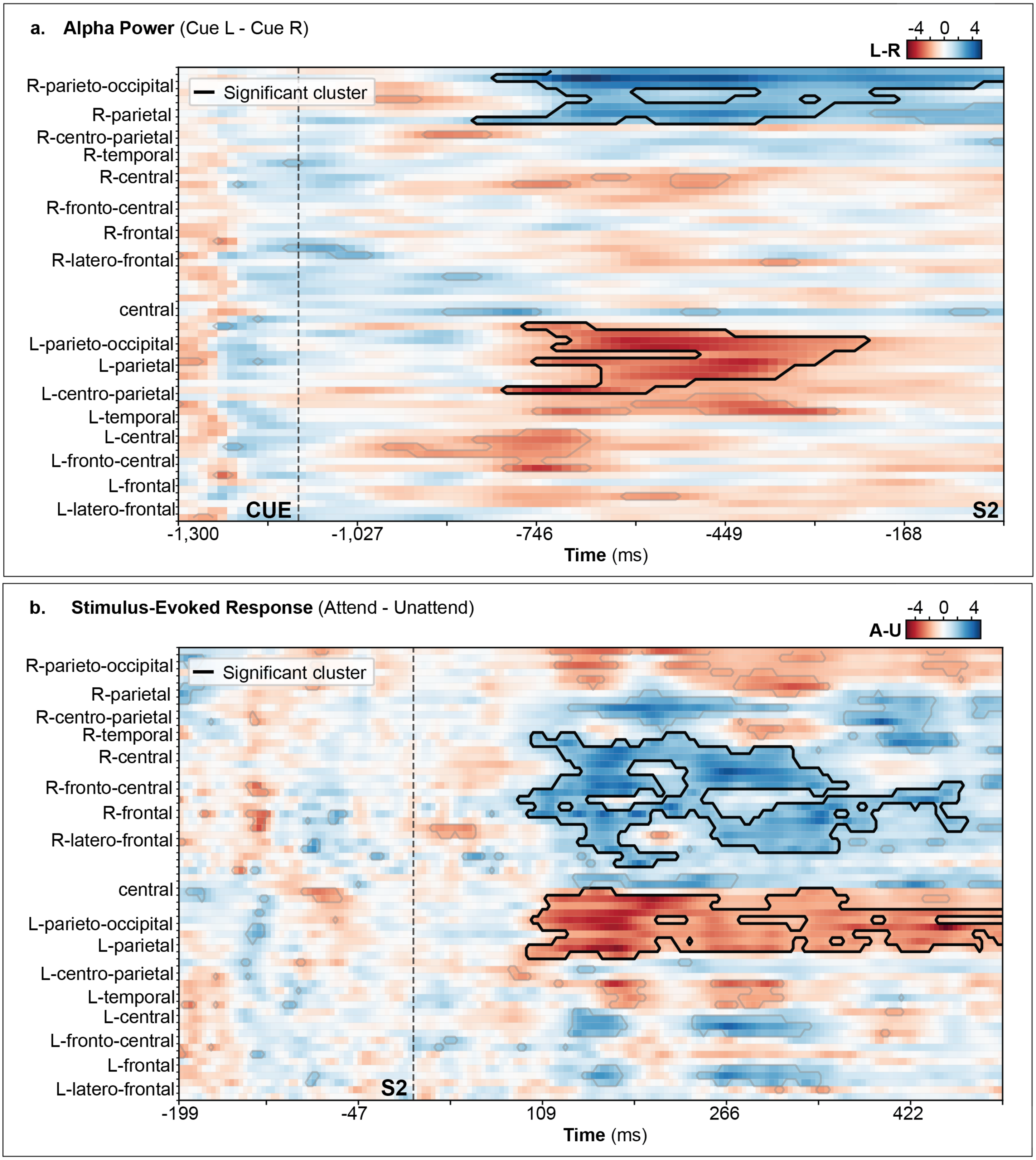


**Supplementary Figure 1. Spatiotemporal cluster-based permutation analysis.**

**a.** Anticipatory alpha power was assessed using a one-sample, cluster-based permutation *t*-test comparing cue-left versus cue-right trials (Cue L − Cue R) across sensors and time during the pre-stimulus interval. Data were collapsed across pure and inter-mixed blocks. The heatmap shows *t*-values over time (−1300 to 0 ms relative to S2 onset). Black contours indicate spatiotemporal clusters that survived permutation correction for multiple comparisons (cluster-level $p < 0.05$). The dashed vertical line denotes cue onset. Topographical maps depict Cue L − Cue R alpha power distributions in successive 50-ms windows, demonstrating maximal lateralized modulation over posterior ROIs. Red indicates greater alpha power for Cue L versus Cue R; blue the converse.

**b.** Evoked responses to unilateral stimuli were assessed using a one-sample, cluster-based permutation *t*-test comparing attended versus unattended conditions (Attend − Unattend) across sensors and time. Attended trials correspond to stimuli presented in the cued hemifield, whereas unattended trials correspond to stimuli in the uncued hemifield. Data were collapsed across pure and inter-mixed blocks. The heatmap shows *t*-values over time (−200 to 500 ms relative to S2 onset). Black contours indicate spatiotemporal clusters that survived permutation correction for multiple comparisons (cluster-level $p < 0.05$). The dashed vertical line at t=0 ms denotes stimulus onset. Topographical maps depict Attend − Unattend scalp distributions in successive 50-ms windows, demonstrating maximal attentional modulation over posterior ROIs.

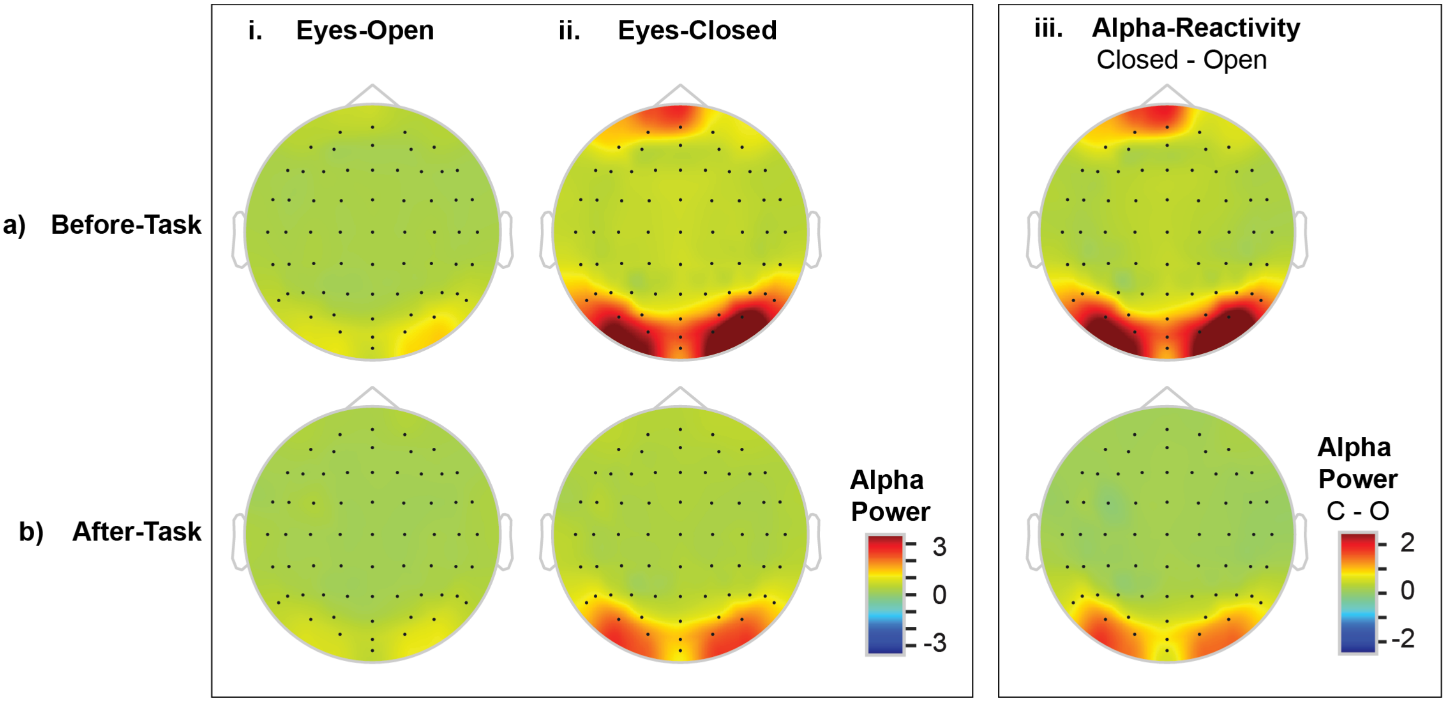


**Supplementary Figure 2. No hemispheric differences in alpha power at rest.**

Topography of total alpha-band (7-13-Hz) power for 2-minutes of resting data for eyes-open (left) and eyes-closed (middle) conditions. Alpha reactivity, measured as the difference between eyes-closed minus eyes-open alpha power is shown on the rightmost subplot. Topographies shown separately for before-task (top) and after-task (bottom) collection of resting state data.

**FRONTAL THETA PHASE MODULATES ASYMMETRIC POSTERIOR NEURAL MECHANISMS OF SPATIAL ATTENTION**
Darrell M, Vanneau T, Brittenham C, Foxe JJ, Molholm S

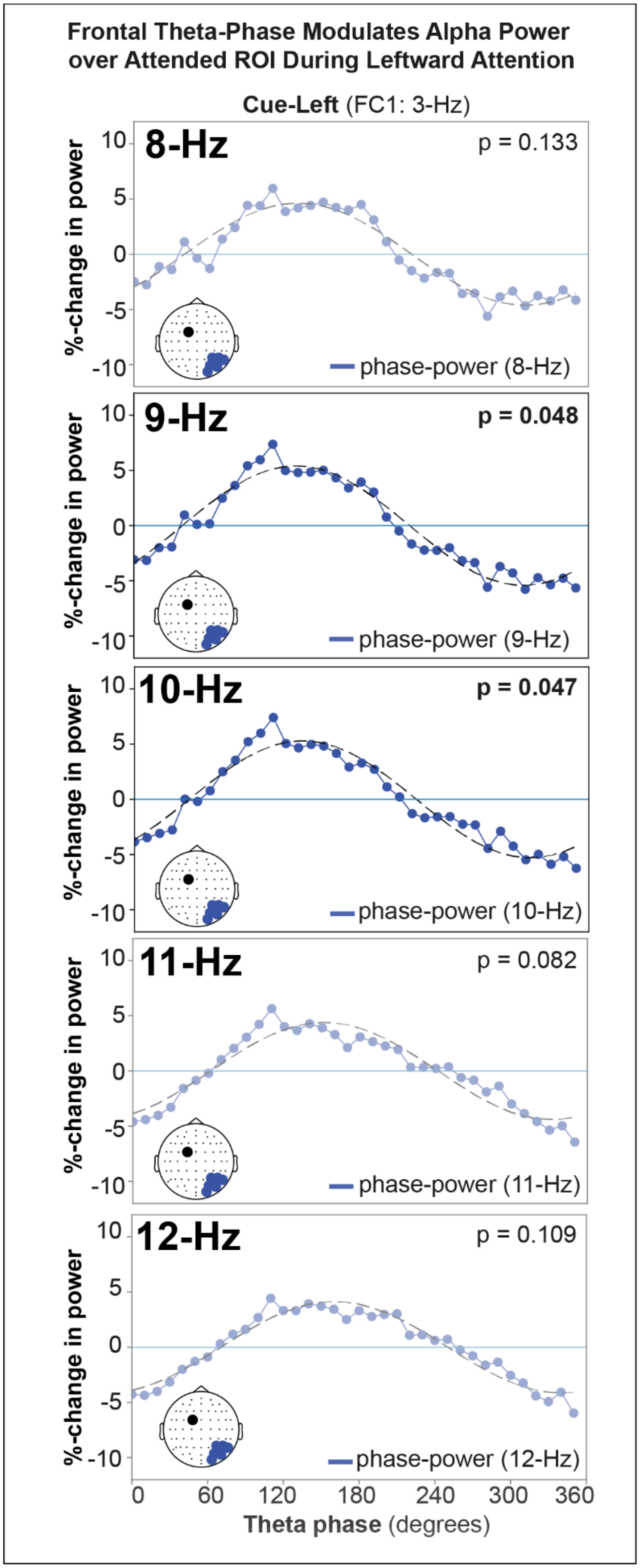

**Supplementary Figure 3. Ipsilateral-cue frontal theta phase–dependent modulation of parieto-occipital alpha power over the attended ROI in leftward-directed attention.**

Theta phase–power functions with phase extracted from fronto-central electrodes at the frequencies identified in Fig. 6a for cue-left trials (3-Hz at FC1). Power at each frequency within the alpha band was measured from parieto-occipital channels contralateral to the to-be-attended location (right parieto-occipital for cue-left) at the same time points used for instantaneous phase extraction. Blue traces show mean percent change in power across theta phase bins (0–360°), with negative values indicating reduced power. Black dashed lines denote the best-fitting one-cycle FFT. Reported p-values reflect the significance of the one-cycle phase–power FFT amplitude derived from permutation testing, indicating a phase-dependent modulation of power. Correlations between theta-phase and 7-Hz ($p = 0.756$) and 13-Hz ($p = 0.086$) were omitted from the figure due to space constraints.

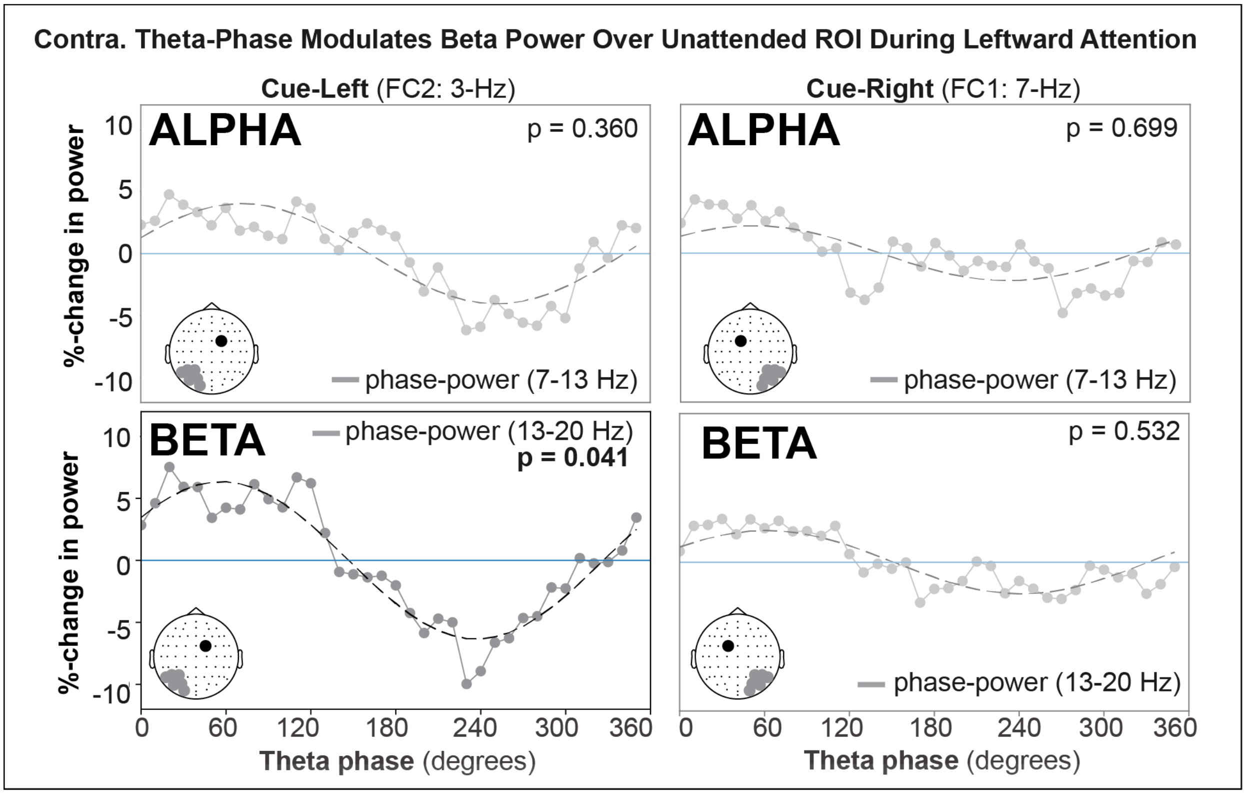


**Supplementary Figure 4. Contralateral-cue frontal theta phase–dependent modulation of parieto-occipital neuro-oscillatory power over unattended ROI.**

Theta phase–power functions with phase extracted at the frequencies identified in Fig. 6a, from contralateral fronto-central electrodes: cue-left trials at FC2 (3-Hz; left) and cue-right trials at FC1 (7-Hz, right). Although a significant phase–behavior relationship was also detected at 6-Hz over FC2, we present only the 7-Hz results for clarity and brevity, as the 6-Hz effect demonstrated an equivalent phase–power coupling profile at this site.

Power was measured from parieto-occipital channels contralateral to the to-be-ignored (unattended) location (left parieto-occipital for cue-left; right parieto-occipital for cue-right) at the same time points used for instantaneous phase extraction for alpha- (7-13-Hz; top) and beta- (13-20 Hz; bottom) bands. Gray traces show mean percent change in power across theta phase bins (0–360°), with negative values indicating reduced power. Black dashed lines denote the best-fitting one-cycle FFT. Reported p-values reflect the significance of the one-cycle phase–power FFT amplitude derived from permutation testing, indicating a phase-dependent modulation of power.

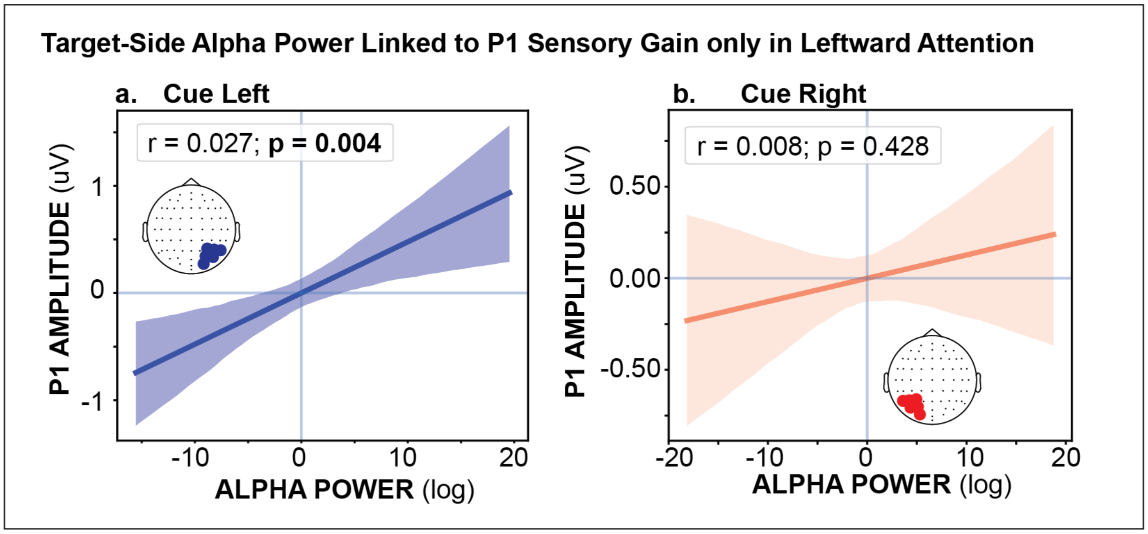


**Supplementary Figure 5. Preparatory parieto-occipital neuro-oscillatory power modulates sensory gain only in leftward attention.**

Correlations between preparatory alpha power and P1 amplitude over the attended ROI (cue-left: right hemisphere; cue-right: left hemisphere) for cue-left **(a)** and cue-right **(b)** trials. Preparatory alpha power was computed from parieto-occipital electrodes during the −800 to −200 ms pre-stimulus interval, indexing anticipatory sensory gating, and P1 amplitude was extracted from the 110–150 ms post-stimulus window, reflecting early sensory gain of the evoked response. Only trials containing attended stimuli were included to ensure that P1 measurements corresponded to attended processing.

| | **Hemisphere Effects** | **Condition (Eyes-Open vs. Eyes-Closed) Effects** | | **RS-Block Effects** | **Age Effects** |
|---|---|---|---|---|---|
| | | **Condition Effect** | **Condition: Interaction between Condition and RS-Block** | | |
| **Resting alpha power** (Morlet) | *None observed* | Main effect of Condition; driven by greater alpha power during eyes-closed (vs. eyes-open) state | Condition x RS-Block; driven by greater alpha power during eyes-closed (vs. eyes-open) state. Effect modulated by RS-Block, such that the eyes-closed power was higher before-task (vs. after-task) with no difference in eyes-open power, resulting in a greater eyes-closed vs. -open difference in the before-task resting state | Main effect of RS-Block duration; driven by greater alpha power before task (vs. after task) | *None observed* |
| **Resting alpha frequency** (FOOOF) | *None observed* | *None observed* | *None observed* | Main effect of RS-Block duration; driven by higher alpha frequency before task (vs. after task) | Main effect of Age; driven by higher peak alpha frequency in older subjects |
| **Aperiodic exponent** (FOOOF) | *None observed* | Main effect of Condition; driven by greater aperiodic exponent during eyes-closed (vs. eyes-open) state | *None observed* | *None observed* | Main effect of Age; driven by lower exponents in older subjects |

**Supplementary Table 1: Resting alpha dynamics.** Summary of LMM results for resting alpha power, frequency and aperiodic exponents in parieto-occipital ROI. LMM included fixed effects of Hemisphere (left/right parieto-occipital ROI) × Condition (eyes-open/eyes-closed) × RS-Block (before-task/after-task), with Age entered as a covariate, and a random intercept for Subject *(1 | Subject)*.
